# Parallel Visual Circuitry in a Basal Chordate

**DOI:** 10.1101/514422

**Authors:** Matthew J. Kourakis, Cezar Borba, Angela Zhang, Erin Newman-Smith, Priscilla Salas, B. Manjunath, William C. Smith

**Author notes:** These two authors contributed equally to this work (Matthew J. Kourakis, Cezar Borba).

## Abstract

A common CNS architecture is observed in all chordates, from vertebrates to basal chordates like the ascidian *Ciona*. Currently *Ciona* stands apart among chordates in having a complete larval CNS connectome. Starting with visuomotor circuits predicted by the *Ciona* connectome, we used expression maps of neurotransmitter use with behavioral assays and pharmacology to identify two parallel visuomotor circuits that are responsive to different components of visual stimuli. The first circuit is characterized by glutamatergic photoreceptors and responds to the direction of light. These photoreceptors project to cholinergic motor neurons, via two tiers of cholinergic interneurons. The second circuit is responsive to changes in ambient light and mediates an escape response. This circuit starts with novel GABAergic photoreceptors which project to GABAergic interneurons, and then to cholinergic interneurons shared with the first circuit. Our observations on neurotransmitter use and the behavior of larvae lacking photoreceptors indicate the second circuit is disinhibitory.

---

Until the recent description of the connectome of a larva of the acidian *Ciona* [1], whole-nervous system connectomics was limited to the nematode *C. elegans*, although extensive connectomes are now available for *Platynereis* [2] and *Drosophila* [3, 4] larvae. Both the small scale of synapses (< 0.5 μm), which necessitates serial section electron microscopy (ssEM) or other similar methods, and the inherent difficulty in visualizing and tracing synaptic connections has so far limited the number of connectomic descriptions of complex nervous systems to discrete CNS regions [5, 6]. Even for the simplest vertebrate models, such as larval zebrafish with ~100,000 neurons [7], a full connectome is likely many years away. Given the challenges of vertebrate connectomics, the ascidian larva provides a unique model for bridging connectomic-level understanding of simple invertebrate models and vertebrates [1]. Ascidians, including those of the widely studied ascidian genus *Ciona*, are members of the chordate sub-phylum Tunicata, which comprise the closest extant relatives of the vertebrates [8]. While much smaller and containing many fewer cells, ascidian tadpole larvae resemble vertebrate larvae in having a prominent notochord running the length of a muscular tail and a dorsal CNS with regional anterior-to-posterior homology to vertebrate CNSs (see recent review, [9]). Despite these homologies with vertebrates the *Ciona* CNS contains only 177 neurons [1]. The *Ciona* larval CNS consists of the *brain vesicle* (BV; also known as the *sensory vesicle*), a region homologous to the vertebrate forebrain and midbrain, followed by the *neck* region, a homolog of the vertebrate midbrain/hindbrain junction (Figure 1). Immediately posterior to the neck is the *motor ganglion* (MG; also known as the *visceral ganglion*). The MG is thought to be homologous to the vertebrate hindbrain, and contains ten motor neurons (MN) as well as number of interneurons - including the two *descending decussating neurons* (ddN) which have been equated with vertebrate Mauthner cells which mediate the startle response [10]. The *Ciona* PNS consists of 54 neurons [11, 12], including several neural crest-derived peripheral neurons [13], and several classes of mechanosensitive (and possibly chemosensitive) peripheral epidermal nerves [12].

Ascidians such as *Ciona* have a biphasic lifecycle. Ascidians spend their first few days as freely swimming tadpole larvae. The larvae then attach to a substrate and undergo metamorphosis to form a sessile adult [14]. As larvae, ascidians are tasked with locating a suitable settling substrate while avoiding predation. Among the described behaviors of *Ciona* larvae are geotaxis, a mechanosensory/touch response, an escape response mediated by light dimming, and negative phototaxis [15–18]. Several sensory systems are found in the BV, including the geotactic otolith and the photosensitive ocellus. Also found in the BV are the coronet cells, an enigmatic group of sixteen dopamine-producing cells [19] that appear to be sensory neurons [1], but of unknown function. Input from sensory neurons of the BV and PNS is directed to the MG, in most cases via one or two interneurons [1, 12].

The *Ciona* connectome provides a detailed and quantitative connectivity matrix of the 6618 chemical and 1206 electrical CNS synapses in a *Ciona* larva [1] and serves as a framework from which neural circuits can be predicted and investigated. Figure 1 shows a simplified visuomotor circuit proposed from the connectome data [1]. In this figure, neurons of the same type are clustered (e.g., photoreceptors) and the number of neurons of each type is indicated in parentheses. The full connectivity for the visuomotor circuit showing all neurons along with chemical and gap junction/electrical synapses (and their relative strengths) is shown in Figure S1a and b, respectively (derived from data tables in [1]). As shown in Figure S1b, gap junctions are few and relatively small in the BV and become more prominent in the MG.

The primary photoreceptive organ of *Ciona* is the ocellus, which consists of two groups of photoreceptors, three lens cells, and one pigment cell. The first photoreceptor group (PR-I) is comprised of 23 cells and clustered around the pigment cell (Figure 1). The opsin-containing outer segments of the PR-Is project into the cup-shaped pigment cell making this group sensitive to the direction of incident light, thereby mediating negative phototaxis [17, 20]. The second group of ocellus photoreceptors (PR-II, Figure 1) contains seven cells and is adjacent and anterior to the PR-Is, and is not associated with the pigment cell. The PR-II cluster mediates the light dimming response by evoking highly tortuous and leftward-biased swims [17]. There is a third set of six photoreceptors (PR-IIIs) distal to the ocellus of unknown function, although they are not involved in either phototaxis or the dimming response [20].

The PR-I and -II photoreceptors synapse primarily onto two classes of relay neurons (RNs) in the posterior BV (pBV). The six *photoreceptor RNs* (prRN) receive input exclusively from the PR-I photoreceptors and then project posteriorly to the paired right/left *MG interneurons* (three on each side; MGIN in Figure 1 and SFigure 1a). A second cluster of eight RNs, the *photoreceptor ascending MG RNs* (pr-AMG RN) are postsynaptic to both the PR-I and PR-II photoreceptors, and are so-named because they, unlike the prRNs, receive input from the *ascending MG peripheral interneurons* (AMG neurons; not shown in Figure 1). There are also extensive synaptic connections between the pr-AMG RNs and the prRNs. Like the prRNs, the pr-AMG RNs project posteriorly to the left and right MGINs. The MGINs in turn synapse onto the paired right and left motor neurons (five on each side).

The *Ciona* connectome thus predicts a complete visuomotor circuit from photoreceptors to muscle target cells, and provides a valuable comparative model to their close cousins the vertebrates, as well to other well-described but much more distantly related models such as *Drosophila, C. elegans* and *Platynereis*. In our previous work we presented results showing that the PR-I photoreceptors are sensitive to the direction of light while the PR-II photoreceptors are sensitive to changes in the intensity of ambient light [17]. In the current study we find that neurotransmitter use in visuomotor circuits, as well as behavioral assays and pharmacology, indicates two distinct pathways. The direction of light is detected by glutamatergic photoreceptors that evoke an excitatory pathway. In contrast, the intensity of light is detected by a second pathway containing novel GABAergic photoreceptors and interneurons. Both the sequential array of GABAergic neurons in this pathway and the behavior of larvae lacking photoreceptors indicates that this circuit is disinhibitory.

## Results

**Figure 1.**
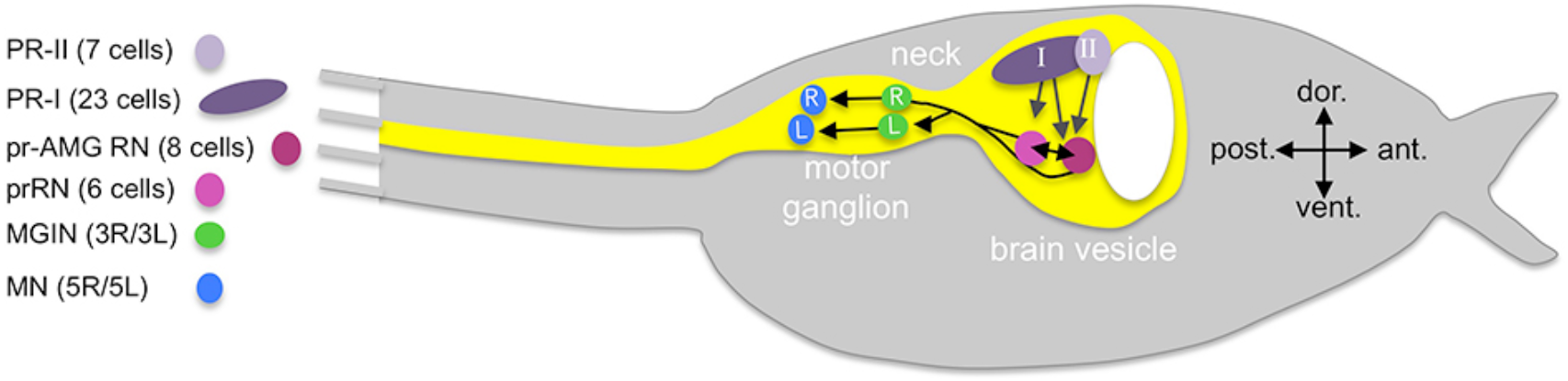
Cartoon of Ciona tadpole larva with outline of central nervous system (yellow). The minimal visuomotor circuit is shown with circles representing classes of neurons with the number of cells of each class indicated in the parentheses of the key. Abbreviations: dors., dorsal; vent., ventral; anter., anterior; post., posterior; PR-II, photoreceptor group II; PR-I photoreceptor group I; pr-AMG RN, photoreceptor ascending motor ganglion relay neuron; prRN, photoreceptor relay neuron; MGIN, motor ganglion interneuron; MN, motor neuron. L, left; R, right. Cell types are color coded according to [1].

### Glutamatergic and GABAergic Photoreceptors

The *Ciona* connectome provides a detailed description of chemical synapse connectivity but it provides no information on neurotransmitter (NT) use. While the expression of genes in the *Ciona* CNS and PNS that mark NT use [e.g., vesicular glutamate transporter (VGLUT), vesicular GABA transporter (VGAT), tyrosine hydroxylase (TH), and vesicular acetylcholine transporter (VACHT) for glutamatergic, GABAergic/glycinergic, dopaminergic and cholinergic neurons, respectively] has been extensively reported [19, 21–24], finding exact matches of expression patterns to neurons, or group of neurons, in the connectome is not always possible. For example, the ocellus is reported to have widespread VGLUT expression, indicating that the *Ciona* photoreceptors, like those of vertebrates are glutamatergic [22]. However, the expression domains of both VGAT and glutamic acid decarboxylase (GAD) are suggestive of a subpopulation of GABAergic/glycinergic photoreceptors [25, 26], although the identies of these cell within the ocellus is not known. To investigate this further, fertilized eggs from a stable transgenic *Ciona* line expressing *kaede* fluorescent protein under the VGAT promoter (pVGAT>kaede) [27] were microinjected with a pOpsin1>red fluorescent protein (RFP) construct [28, 29] (Figure 2a). We observed a subset of photoreceptors coexpressing the two fluorescent markers both at the anterior and ventral sides of the ocellus (white and orange arrows, respectively). In the field of view shown in Figure 2a the eminens cells is also evident due to its expression of VGAT (white arrow), in agreement with earlier reports with GAD [24]. To investigate BV VGLUT and VGAT expression in greater detail we used Hybridization Chain Reaction *in situ* (HCR *in situ*) [30] (Figure 2b). In agreement with previous reports [22] we observed VGLUT expression in the ocellus (blue arrowhead), the two otolith antenna cells (AC), and in epidermal sensory neurons (red arrowheads). Consistent with the above transgenic data, VGAT was expressed in two separate clusters within the ocellus (white and orange arrowheads), as well as in a separate group of BV neurons outside the ocellus corresponding to previously described VGAT-positive neurons that project axons to the MG [25] (labeled in Figure 2b as RNs, see next section). The anterior VGAT-expressing photoreceptor cluster consisted of 7 cells (±1 cell, n=7 larvae), while the posterior group consisted of two cells in all samples. We also observed a subset of the VGAT-expressing cells in both the anterior and posterior clusters that also expressed VGLUT (Figure 2c and d). In the anterior cluster, we observed that the 2 cells (± 1; n=4 VGAT/VGLUT double *in situ* larvae) at the anterior edge exclusively expressed VGAT (white arrows in Figure 2c), while the four cells immediately posterior to these cells co-expressed VGAT and VGLUT. In the posterior cluster we observed in all samples that one of two cells coexpressed VGAT and VGLUT, while the other only expressed VGAT (Figure 2c, orange arrows).

**Figure 2.**
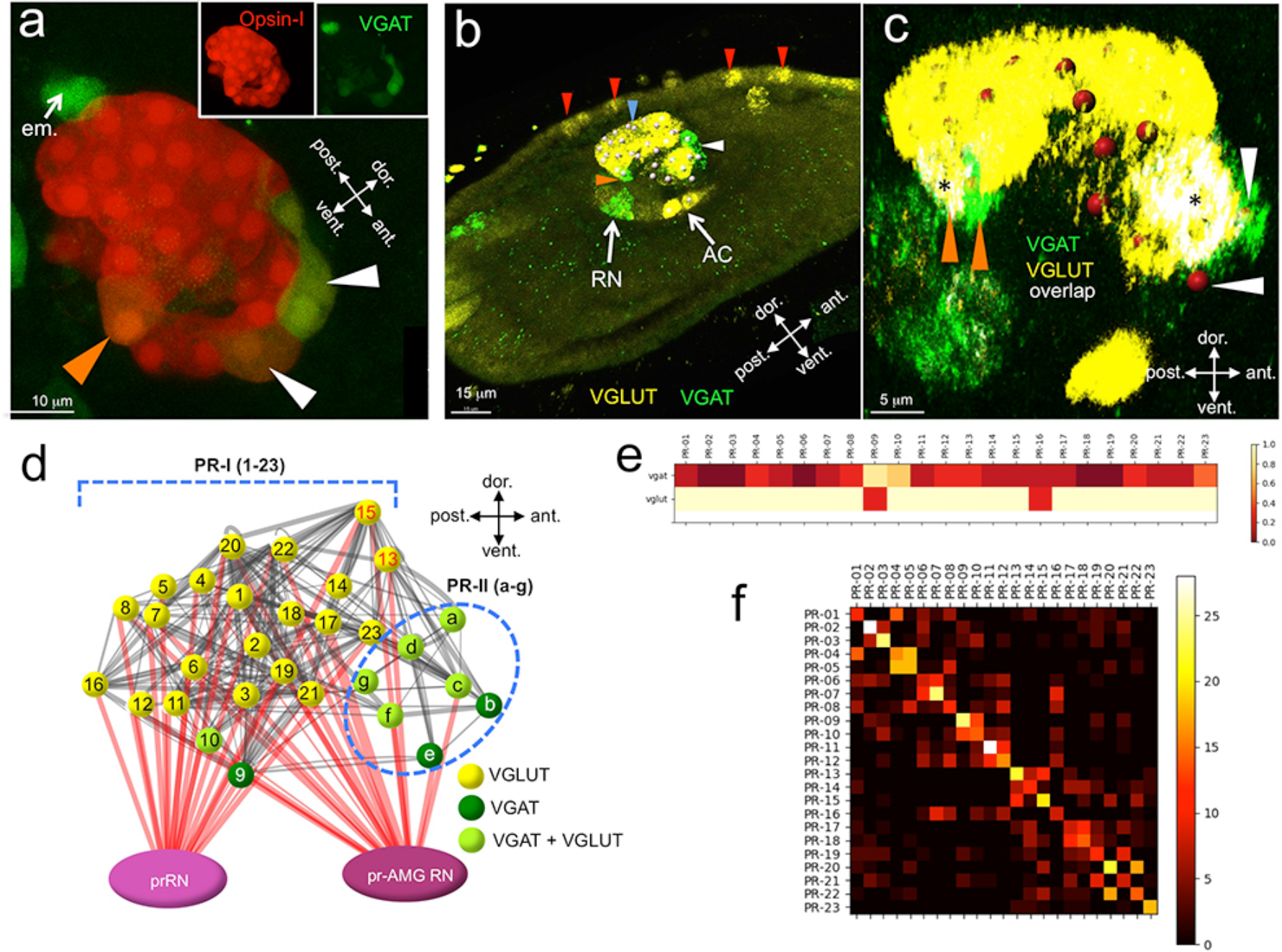
Neurotransmitter use in the ocellus. **a**. Coexpression of opsin and VGAT reporter constructs in the ocellus (white and orange arrowheads). **b**. Expression of VGLUT and VGAT in the brain vesicle and epidermis by in situ hybridization. VGAT was observed in an anterior (white arrowhead) and posterior (orange arrowhead) domain of the ocellus. Blue arrowhead indicates VGLUT expression in the ocellus, and red arrowhead indicate VGLUT-expressing epidermal sensory neurons. **c**. Posterior VGAT-expression in the ocellus consists of two cells (orange arrowheads), one exclusively expressing VGAT, and one coexpressing VGAT and VGLUT. Nuclei are shown as red spheres. **d**. Neurotransmitter predictions color-coded on a schematic diagram of the ocellus photoreceptors. Lines between photoreceptors indicate chemical synaptic connections taken from [1], with red lines indicting projections to the relay neurons. **e**. Heat map for Photoreceptor Group-I neurotransmitter predictions from registration. **f**. Confusion matrix of cell registration. High values (light colors) in the diagonal indicate higher confidence. Abbreviations: dors., dorsal; vent, ventral; ant., anterior; post., posterior; em., eminens cell; RN, relay neuron; AC, antenna cells; pr-AMG RN, photoreceptor ascending motor ganglion relay neuron; prRN, photoreceptor relay neuron; VGAT, vesicular GABA transporter; VGLUT, vesicular glutamate transporter.

Given the cellular anatomy of the *Ciona* ocellus, with seven PR-IIs anterior and 23 PR-Is posterior [1, 20], and our transgenic and HCR *in situ* results, we assign the PR-IIs as being VGAT-positive, with a subset co-expressing VGLUT (Figure 2d). The anterior/ventral location of the two VGAT-only PR-IIs suggest that they are PR-b and -e (Figure 2c and d). By contrast, the majority of the PR-Is are exclusively glutamatergic with the exception of two ventral cells, one co-expressing VGAT and VGLUT and the other expressing only VGAT. While the identities of the PR-II subpopulations were evident from the ocellus anatomy, the identities of the two VGAT-expressing PR-Is were initially less clear. To get a better indication of the identities of these two cells we performed a registration of cell centroids from multiple *in situ* datasets (n=11) with the centroids from the connectome ssEM dataset. This registration would only be meaningful if there is strong stereotypy in the number and position of the neurons among *Ciona* larvae. The ocellus photoreceptor soma and their outer segments are known to be arranged in rows, suggesting an ordered cellular architecture [1, 20]. Moreover, we reasoned that stereotypy in the ocellus, if present, should be evident when registering VGAT- and VGLUT-expressing cells, both across multiple *in situ*-stained larvae, and individually to the ssEM photoreceptor centroids. Convergence of NT type with registered photoreceptors (both between HCR *in situ* samples and between these and the ssEM sample) would be taken as evidence of stereotypy, and of the validity of making NT use predictions.

For photoreceptor registration, DAPI-stained ocellus nuclei from VGAT and VGLUT *in situ* labeled larvae were segmented from 3D image stacks to serve as cell centroids (nuclei are indicated in Figure 2c as red spheres). Based on the *in situ* signal, each centroid was designated as VGAT or VGLUT, or both. Finally, the antenna cell and ddN nuclei in the image stacks were segmented to serve as anchor points for registration. Registration of the segmented HCR *in situ* nuclei to each other and to the connectome PR-I nuclei was done according to [31]. Briefly, rotation and affine transformations were applied to each set of *in situ* nuclei coordinates to register them individually to the connectome cell nuclei coordinates. The results are presented as a heat map showing for each PR-I the relative frequencies it registered with an *in situ* centroid of each NT type (Fig 2e). To assess the validity of registration, a confusion matrix was constructed [32] (Figure 2f; Supplemental Table 1). In this analysis each set of HCR *in situ* centroids was registered to all other HCR *in situ* datasets, and to the connectome centroids. The confusion matrix shows the number of times a registration of the HCR *in situ* centroid to a connectome centroid corresponds with the registration of another HCR *in situ* dataset. The higher the values along the diagonal of the matrix, the more the datasets agree with each other as to registration to the connectome centroids. From the matrix we observed strong overall support for the registration, although with variable confidence for each photoreceptor. The heat map indicates that among the PR-Is, PR-9 is likely to be exclusively VGAT-positive, while PR-10 is likely to be both VGAT- and VGLUT-positive. The confusion matrix gives high confidence to this assignment, particularly for PR-9. Although PR-9 and PR-10 appear to stand out from the other PR-Is in their NT use, the connectivity of these two photoreceptors in the visuomotor pathway does not appear to be qualitatively different than the other PR-Is (Figure S2). Finally, the heat map confirms that the other PR-Is are exclusively VGLUT, however, with lower confidence for PR-16, which failed to register well.

### Posterior Brain Vesicle Relay Neurons are mixed VGAT and VACHT expressing

Sensory input from the photoreceptors, antennae cells, coronet cells, bipolar tail neurons and a subset of peripheral neurons is directed to a cluster of ~30 RNs in the pBV. These RNs in turn extend axons through the neck to the MG. Among this cluster are the six prRNs and eight pr-AMG RNs (Figure 1; [1]). Previous in situ hybridization studies identified VGAT- and VACHT-expressing neurons in appropriate place in the BV to be RNs [25]. Moreover, these neurons project axons proteriorly to the MG, as is definitive of the pBV RNs. BV neurons expressing other major NTs, including glutamate, dopamine, and serotonin, are neither in the correct brain region to be RNs, nor do they project from the BV to the MG ([19, 22, 23], and our observations). By HCR *in situ* we observed that the pBV RNs cluster in two distinct groups in the anterior/posterior axis, with the anterior cluster expressing VACHT, and the posterior group expressing VGAT (Figure 3a). We observed an average of 16 (± 1.6, n=9 larvae) VGAT-positive neurons and 11 (± 1, n=8 larvae).

Unlike the ocellus, the pBV RN cluster does not have obvious anatomical features, although the various classes of RNs are clustered, with, for example, the antenna cell RNs being posterior to the photoreceptor RNs (Figure S3; [1]). However, given the diversity of RN types in the pBV it is unlikely that the expression domains of VGAT and VACHT precisely correspond to the clusters of RN classes. In order to make predictions of NT use in the RNs we used the same registration approach as with the photoreceptors (n=7 VGAT/VACHT double *in situ* datasets, Figure 3S). The confusion matrix for the RNs shows a lower level of convergence in the registration than for the PR-Is, suggesting that the cellular anatomy of the RN cluster is less structured than the ocellus (Figure 3b; Figure S3; Supplemental Table 2). However the confusion matrix also shows that in the registration the RNs are most often confused for other RNs of the same class (white boxes in 3b). This is most evident when the registration is done not with single cells, but with pooled RNs of each classes (Figure 3c; Supplemental Table 3), and is presumably a reflection of the clustering of RN classes in the pBV. Thus we can have higher confidence in the NT use by RN class than we can have in individual neuron identities. For example, the connectome shows the antenna RNs (AntRN) are clustered at the rear of the BV (Figure S3; [1]), as are the VGAT expressing neurons (Figure 3a; Figure S3). Accordingly, the registration predicts that eight of the ten AntRNs are VGAT positive (Figure 3c). For the present study, which focuses on the visuomotor pathway, the registration predicts that five of the eight pr-AMG RNs are VGAT expressing, two are VACHT expressing, and one (pr-AMG RN 157) cannot be resolved (no dual VGAT/VACHT expression was observed in the *in situs*). On the other hand, the registration predicts that the six prRNs are evenly mixed between VGAT and VACHT expression. These predictions provide starting points for experimental validation detailed below.

**Figure 3.**
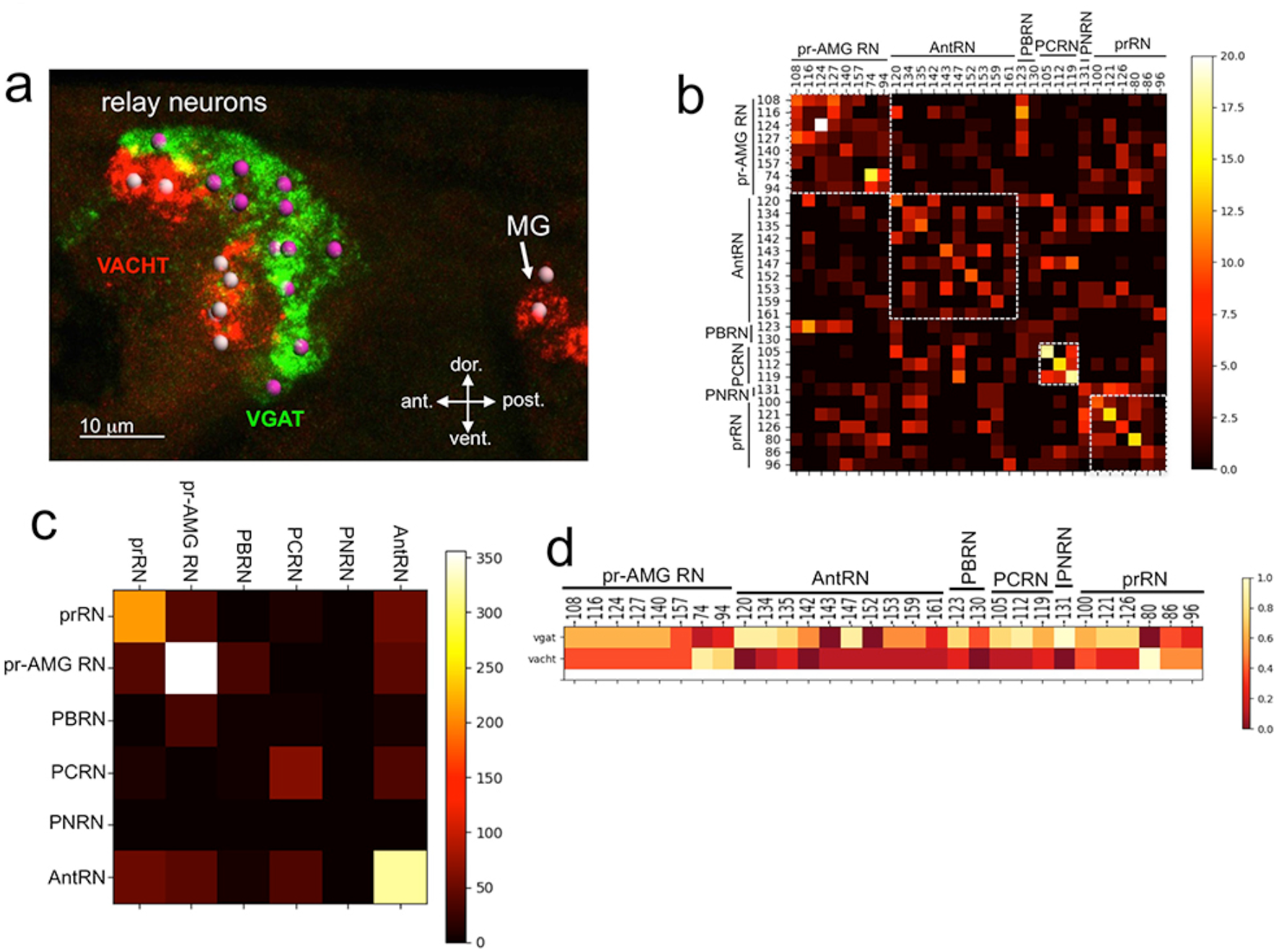
Neurotransmitter use in the relay neurons. a. In situ hybridization of VGAT and VACHT to the relay neurons in the brain vesicle. Also visible is the anterior tip of the motor ganglion. Nuclei are shown as circles. b. Confusion matrix for relay neuron registration. c. Confusion matrix for relay neurons grouped by type. d. Heat map of neurotransmitter predictions from cell registration of relay neurons. Abbreviations: ant., anterior; post., posterior; RN, relay neuron; ant., antenna cells; pr-AMG RN, photoreceptor ascending motor ganglion relay neuron; prRN, photoreceptor relay neuron; AntRN, antenna cell relay neuron; PBRN, photoreceptor-bipolar tail neuron relay neuron; PCRN, photoreceptor-coronet relay neuron; PNRN, peripheral relay neuron; VGAT, vesicular GABA transporter; VACHT, vesicular acetylcholine transporter.

### The motor ganglion contains a mixture of cholinergic and GABAergic neurons

The MG contains ten MNs and five classes of interneurons: six MGINs, seven AMGs, two ddNs, three *ascending contralateral inhibitory neurons* (ACIN), and two *posterior MG interneurons* [1]. Like the ocellus, the MG has a well defined anterior-to-posterior and dorsal-to-ventral cellular anatomy (Figure 4; [1, 12]). The NT use by some of the MG neurons is already documented, including the motor neurons, which are cholinergic [24, 33]. By HCR *in situ* hybridization we found evidence for VGAT- and VACHT-positive neurons in the MG (Figure 4b), and no VGLUT- or TH-positive cells (data not shown). These results are consistent with previous studies [19, 22]. Likewise it was reported that no serotonergic cells were present in the MG [23]. As with the RNs, the VGAT- and VACHT-expressing neurons in the MG segregated anatomically. We also found a population of 6-7 cells between the AMGs and the MNs (asterisks in Figure 4b), that were not annotated in the connectome as neurons and that failed to label with any of our NT markers. We hypothesize that these are ependymal cells, which are abundant in the nerve cord immediately caudal to this region.

**Figure 4.**
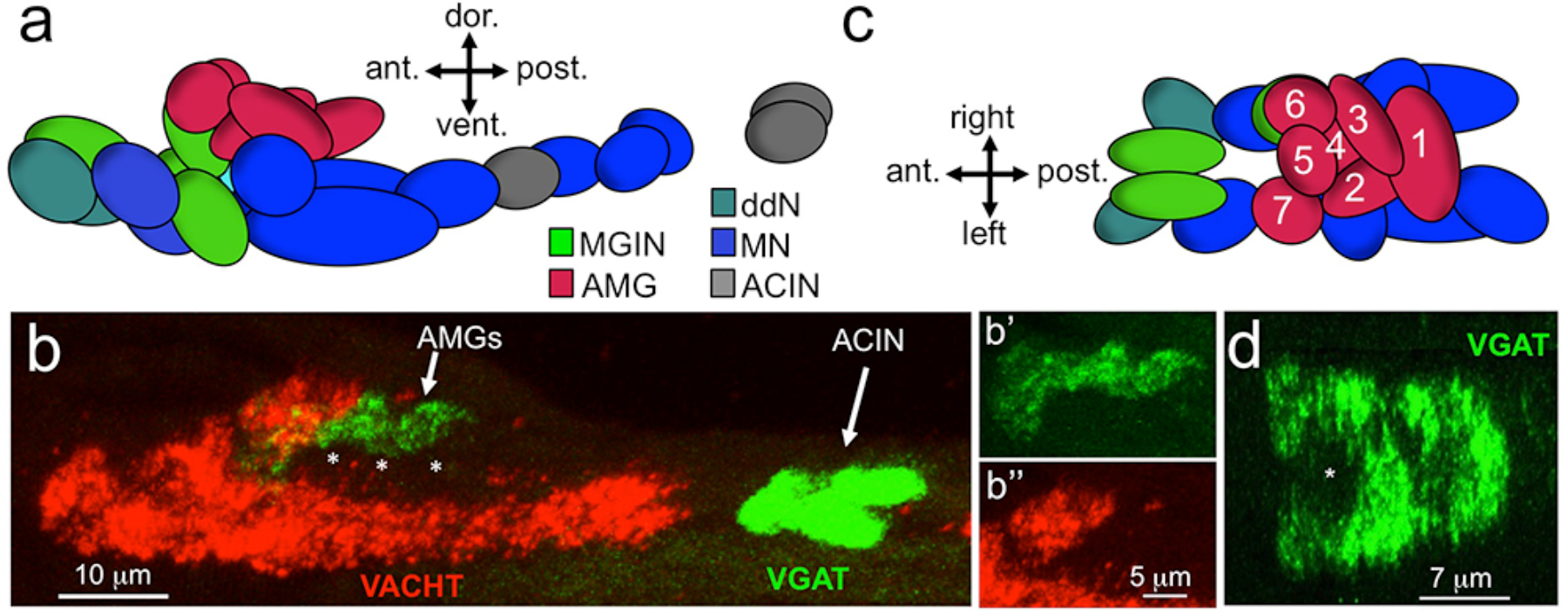
Neurotransmitter use in the motor ganglion. **a**. Diagram of neurons in the motor ganglion (derived from Figure 1 of [10]). Lateral view. **b**. Expression of VGAT and VACHT by in situ hybridization in in the motor ganglion, lateral view. Asterisks indicate predicted ependymal cells. **b’** and **b”** show separate channels for VGAT and VACHT in the AMGs. **c**. Diagram of neurons in the motor ganglion. Dorsal View. AMGs are numbered according to [1]. **d**. Dorsal view of VGAT expression in the motor ganglion. Asterisk indicates central non-VGAT expressing cell. Abbreviations: dors., dorsal; vent., ventral; ant., anterior; post., posterior; AMG, ascending motor ganglion neuron; MGIN, motor ganglion interneuron; ddN, descending decussating neurons; ACIN, ascending contralateral inhibitory neurons; MN, motor neuron; VGAT, vesicular GABA transporter; VACHT, vesicular acetylcholine transporter.

Because of the highly structured MG cellular anatomy (Figure 4a; [1, 12]), we can identify the various MG cell types in the *in situ* data. The anterior group of VGAT-positive cells is clustered dorsally in the MG, and correspond to the AMGs (shown as separate channels in Figure 4b and b). This is most obvious in a dorsal view of the MG (Figure 4c and d), in which a ring of VGAT-positive cells was observed with a central non-VGAT expressing cell in the center (asterisk, Figure 4d). The VGAT-expressing cells appear to be AMGs 1, 2, 3, 4, 6, and 7, while the central cell, which is instead positive for VACHT, appears to be AMG5. Interestingly, the connectome shows that AMG5 differs in its connectivity from the other AMGs. Significantly, AMG5 is the principle synaptic input for PNS neurons. It then synapses to the other AMGs, which in turn project their axons to the pr-AMG RNs in the BV. In the posterior of the MG we observed a pair of VGAT-positive neurons that we can identify as the posterior ACIN pair. Finally, in the ventral MG we observed a continuous block of VACHT expression that encompasses the MNs, ddNs, MGINs, and the anterior ACINs. Identical *in situ* patterns were observed in multiple larvae (Figure S4). For the visuomotor circuit that is the subject of this study (Figure 1), the most important neurons in the MG, the MGINs and the MNs, are all cholinergic.

### Parallel Visuomotor Circuits

Our results indicate that the PR-Is, with the exception of two cells, are glutamatergic, while the PR-IIs are a mixture of GABAergic and GABA/glutamatergic. The *Ciona* genome contains a single glutamate AMPA receptor (AMPAR) [34] that is expressed in larvae in the two antenna cells, and in a small cluster of neurons in the pBV [35]. Published results show that most of the pBV group of AMPAR-positive neurons are clustered at the ends of Arrestin-labeled photoreceptor axons, and that they extend their axons to the MG, suggesting they are photoreceptor RNs (see Figure 2B” in [35]). We find that this pBV group is composed of ~6 cells (Figure S5). To investigate this further, we co-expressed an pAMPAR>GFP construct [35] with pVACHT>CFP and pVGAT>nuclear-RFP constructs. We observed coexpression of the AMPAR reporter in a subset of the VACHT-postive RNs, but never in the VGAT-expressing RNs (Figure 5a).

To assess the function of the AMPAR-positive cells in *Ciona* visuomotor behaviors we used the non-competitive AMPAR antagonist perampanel [36]. For the assay, larvae were treated at 25 hpf with perampanel in sea water and compared to vehicle-treated control larvae for both negative phototaxis and response to light dimming. The negative phototaxis assay consisted of placing the larvae in a 10 cm petri dish of sea water with a 505nm LED lamp placed to one side (described by us previously [17]). Images were collected at 1 minute intervals over 5 hours to assess for taxis (Movie1). Figures 5b and c show representative frames from the time-lapse capture at the start and at 60 min for control and perampanel-treated larvae, respectively. In the control sample the larvae at 60 min were observed to cluster at the side of the petri dish away from the light (distal side; red arrows in Figure 5b). By contrast no taxis was observed in the perampanel treated larvae (Figure 5c). Combined results from three independent assays are shown in Figure 5d and presented as the percent of larvae found on distal third of the petri dish (see Supplemental Data 4). For control larvae ~70% swam to the distal third within 2h, while the perampanel-treated larvae remained evenly distributed across the dish.

The inability of the perampanel-treated larvae to phototax was not the result of an inability to swim, as seen in Movie2 which were taken at 8.9 fps, with and without perampanel. Moreover, we observed that perampanel treatment had no effect on the light dimming response (Movie3). Figures 6a and b show 5-second projection images from Movie3 immediately before and after dimming. In these images swims appear as lines, and the responses in control and perampanel-treated larvae appear qualitatively similar. To quantitatively compare dimming response, control and perampanel-treated larvae were exposed to a range of dimming intensities from 2 to 60-fold and the percentage of larvae responding was measured and presented as a percentage in Figure 6c. The percentage responding at all intensities was very similar for both groups, and pair-wise comparisons at each fold change failed to show significance. In addition, no differences were measured in the velocity or duration of swims in pair-wise comparisons of control and perampanel-treated larvae at any fold-dimming (data not shown). We conclude that there is no change in sensitivity to dimming caused by perampanel treatment, while phototaxis was completely disrupted. Finally, we also observed that the touch response was not inhibited by perampanel (data not shown), despite the presence of VGLUT-positive epidermal sensory neurons [22]. This would appear to agree with the observation that primary RNs for the PNS, the eminens cells and the AMGs do not express the AMPAR ([35]; and our observations). In addition to the AMPAR, the *Ciona* genome contains several other glutamate receptors including one kainate and one NMDA [34], although their expression has not been characterized.

In summary, we are able to separate the phototaxis and dimming behaviors pharmacologically. Moreover, we can identify the VACHT/AMPAR-positive RNs as essential for an excitatory PR-I circuit that involves presynaptic glutamatergic PR-Is and postsynaptic cholinergic MGINs. The number and location of the VACHT/AMPAR-positive RNs, the circuit logic, and our behavioral observations are all consistent with these being prRNs.

**Figure 5.**
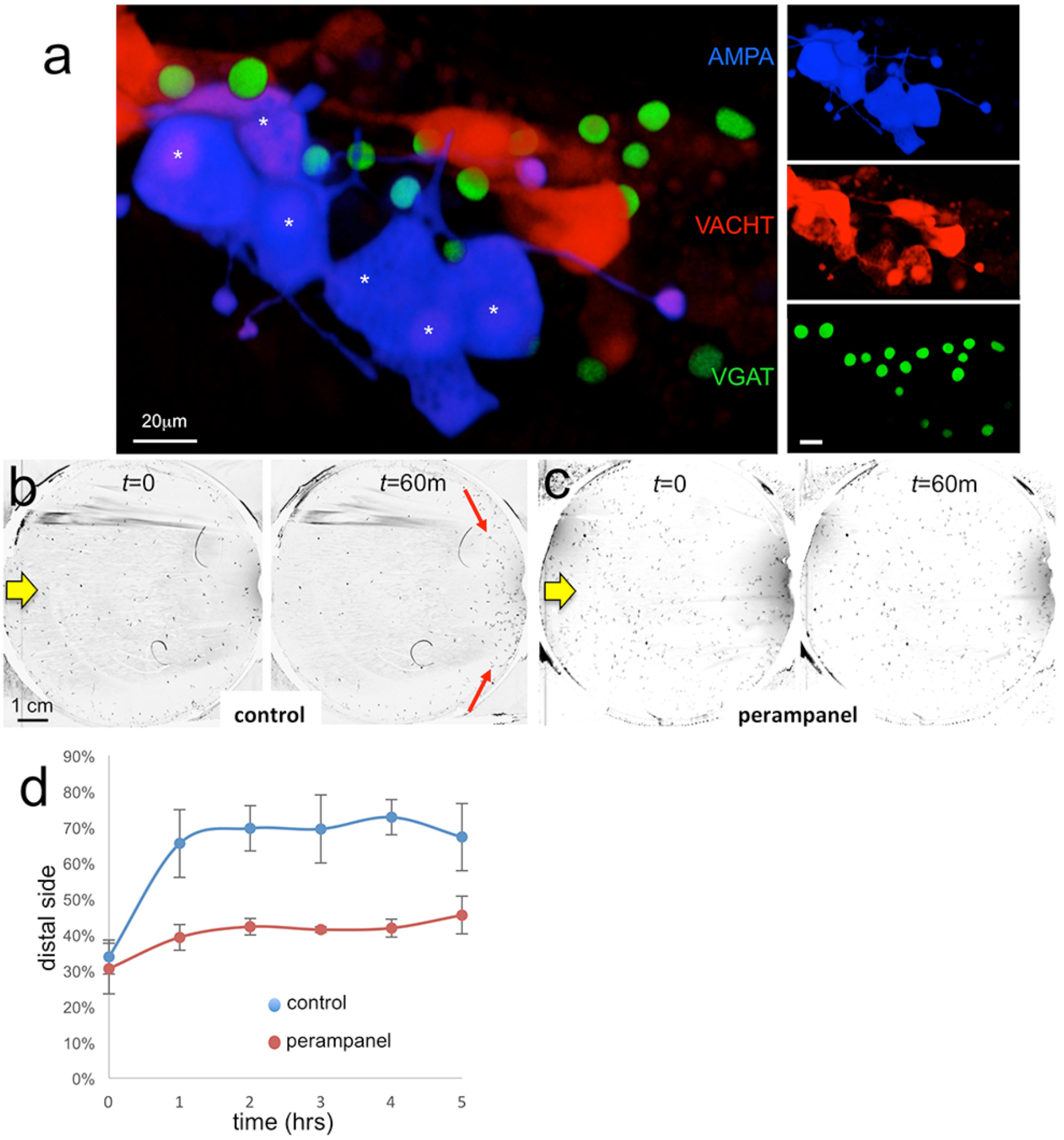
AMPA receptors in negative phototaxis. **a**. Coexpression of an AMPA-receptor and VACHT expression constructs in the relay neurons (white asterisks). The main panel shows the merge while smaller panels at right show single channels. **b**. Negative phototaxis assay in control larvae. Yellow arrow indicates direction of 550nm light. By 60 minutes (m) the majority of the larvae have swum to the side of the dish away from the light (red arrow). **c**. Perampanel treated larvae do not show negative phototaxis. **d**. Quantification of negative phototaxis in control and perampanel treated larvae. Points indicate the averages from three independent assays, + standard deviation. Y-axis represents the percentage of larvae found on the side away from the light source (distal third). Abbreviations: VGAT, vesicular GABA transporter; VACHT, vesicular acetylcholine transporter.

### A disinhibitory circuit

Of equal significance to our observation that navigation is inhibited by perampanel, is our observation that the dimming response, which is mediated by the PR-IIs [17], is not inhibited by perampanel (Figure 6). Our expression studies show that the PR-IIs are comprised of a mixture of VGAT - and VGAT/VGLUT - expressing photoreceptors. Although it is formally possible that PR-IIs signal exclusively via glutamate in an excitatory circuit via a non-AMPA glutamate receptor on their RNs, the fact that several of the PR-IIs are VGAT-only and that the majority of the pr-AMG RNs, the exclusive RNs of the PR-IIs, are predicted to be GABAergic suggests an alternative disinhibitory circuitry logic. This circuit would consist of the inhibitory PR-IIs synapsing to the pr-AMG RNs to reduce their inhibition on the cholinergic MGINs.

**Figure 6.**
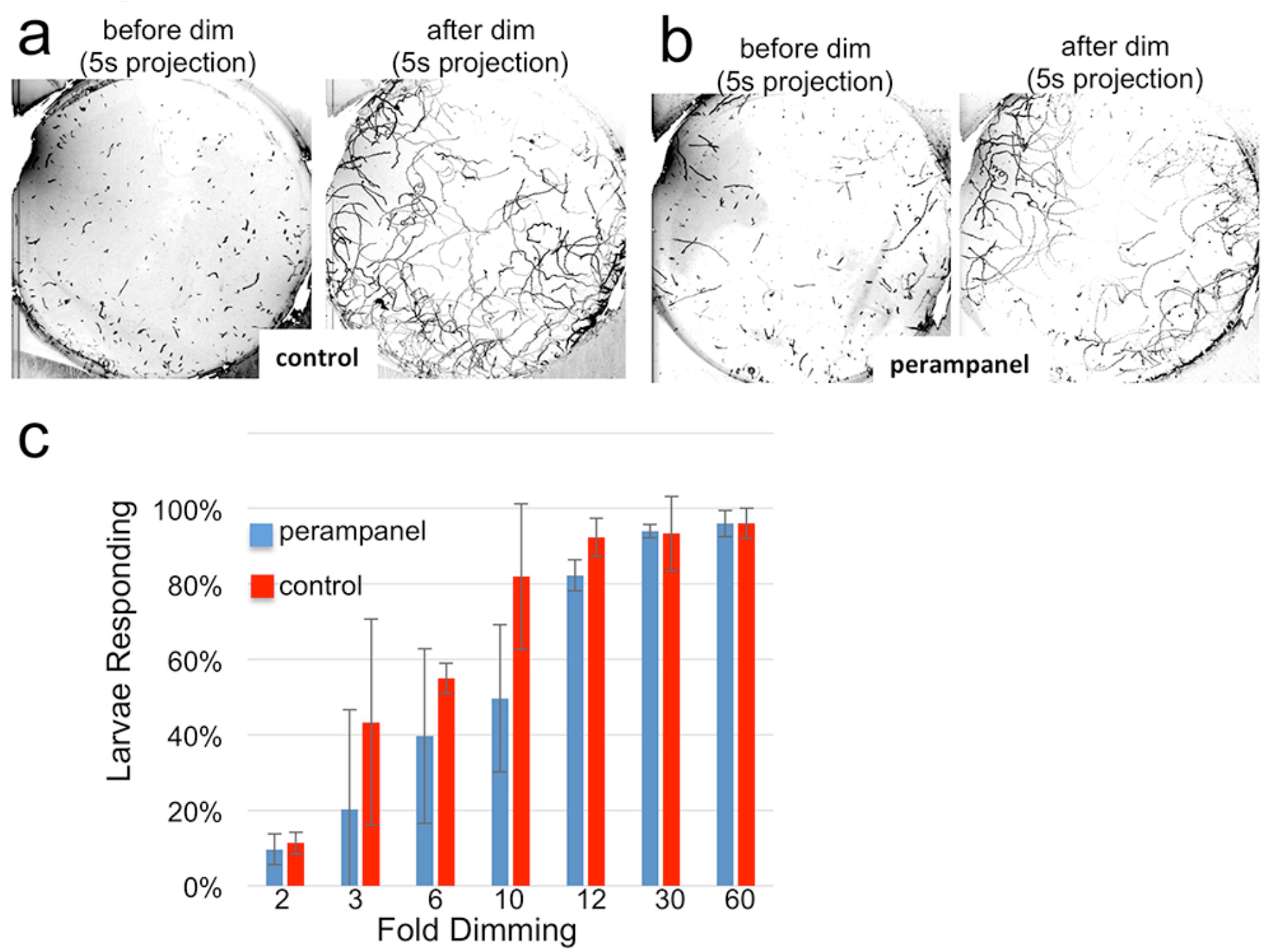
Perampanel does not disrupt the light dimming response. **a**. Light dimming response in control larvae. Shown are 5 second (s) projections from time-lapse movies in which swims appear as lines. Left panel shows a projection 5s before dimming, and right panel 5s after dimming. **b**. same as a, but larvae were perampanel treated. **c**. Quantification of light dimming response in control and perampanel treated larvae. Larvae were exposed to dimming of505nm light from 2- to 60-fold. Dimming response was scored as percent of larvae responding. Bars show averages of three independent assays ± standard deviation.

Implicit in the disinhibitory model is an autonomous level of motor activity in larvae that could be inhibited by the GABAergic pr-AMG RNs, and that this inhibition is released upon stimulation of the GABAergic PR-IIs. To gain evidence for this model we took advantage of a previously described *Ciona* mutant, *frimousse (frm*) [37, 38]. In homozygous *frm* larvae the anterior BV is transfated to epidermis due to a null mutation in a neurula stage-specific connexin gene [38]. *Frm* larvae lack the ocellus pigment cell and photoreceptors, as well as the otolith [37, 38] (Figure 7a). As would be predicted, *frm* larvae show no response to light (our unpublished observation). Despite the defects in the anterior BV, the MG is intact in *frm* larvae as assessed by both gene expression and morphology ([37] and Figure 7a). Moreover not only can *frm* larvae swim, they show increased frequency of spontaneous swims compared to wild type larvae (Figure 7b). In this assay, swimming activity of *frm* and wild type larvae was recorded in 1-min movies at 8.9 fps with 700nm illumination, which the wild type *Ciona* larvae cannot detect [39]. Under these “dark” conditions larval swims consist primarily of randomly occurring short “tail flicks”, with very few sustained swims [17]. Each circle in Figure 7b corresponds to a single larva tracked over one minute, and the number of swim bouts for each larva during the 1-minute is plotted along with the average (red circle) and S.D. (n=75 for both; p< 5×10^-16^). Although the frequency of swims was higher in *frm* larvae, the average swim time was not significantly different between the two (Figure 7c; n=260 and 608, respectively). However a handful of very long swims were observed uniquely in the *frm* group (Figure 7c). Despite the similarity in the swim times between the two groups, the swim characteristics were very different (Movie 4), with the swimming of *frm* larvae being much more stereotyped. For example, while the swim times of wild type and *frm* larvae averaged to very similar values (Figure 7c), the standard deviations of swim times calculated and plotted for each larva show much lower swim-to-swim variation in the *frm* larvae (Figure 7d; p<5 x10^-15^). The stereotypy is even more pronounced when the time interval between swims was analyzed (Figure 7e). For wild type larvae the standard deviations of interval times showed a wide range of values (i.e., high variability of interval times), while the standard deviations for *frm* larvae were much lower (p < 0.0005). The behavior of *frm* larvae is characteristic of an oscillator that evokes spontaneous swims with the frequency of ~8/min. Thus, sensory input from the BV appears to suppress this oscillatory behavior leading to less frequent and more varied swims in wt larvae, supporting a disinhibitory circuit. Interestingly, we observed some VGAT and VACTH expression in remnant of the *frm* BV (Figure 7a). While these expressing cells may be RNs, it remains to be determined whether in the absence of sensory input they develop properly, and if they are functional, how their apparent opposing activities might influence spontaneous swimming (albeit elevated).

**Figure 7.**
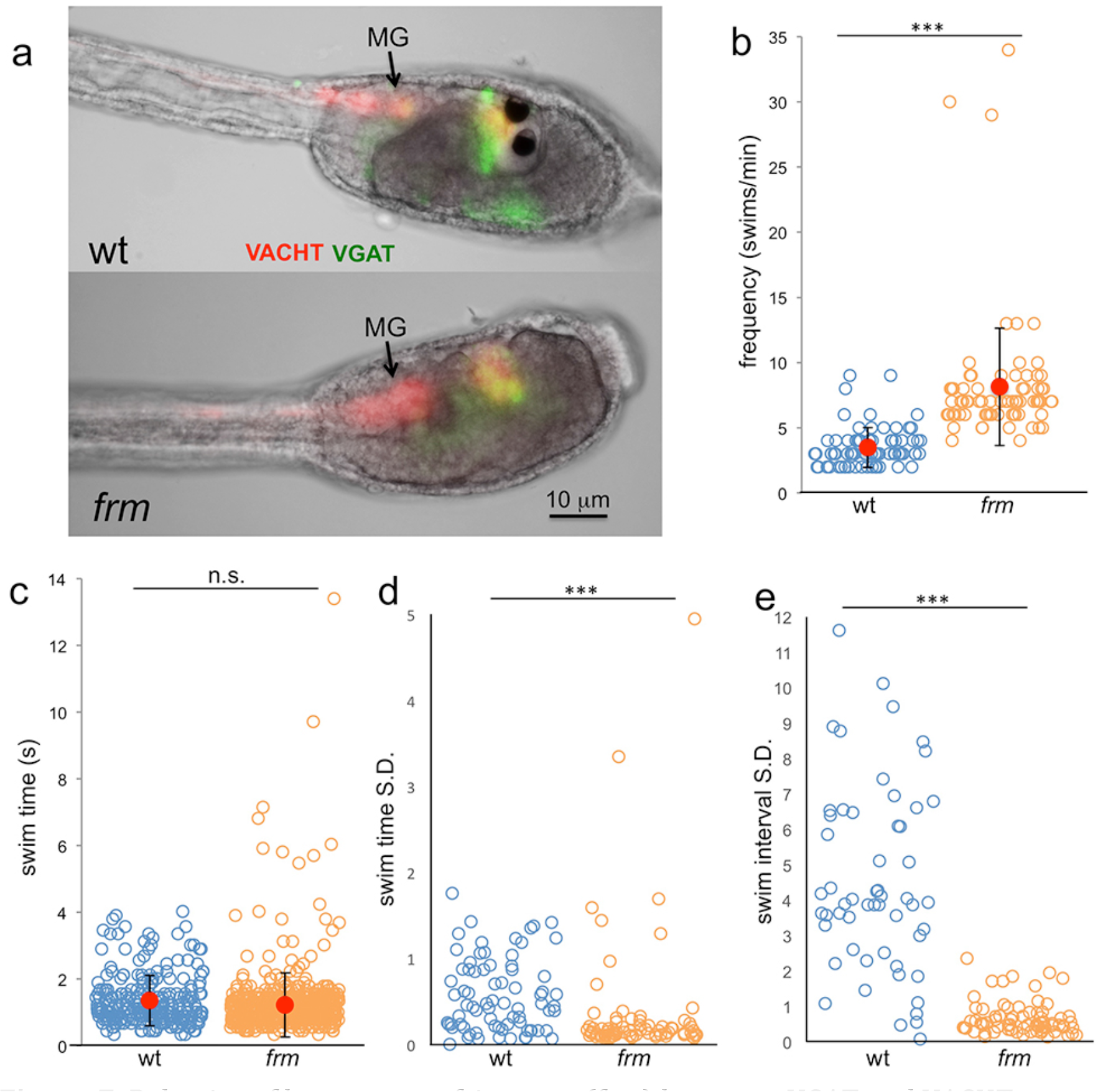
Behavior of homozygous frimousse (frm) larvae. **a**. VGAT and VACHT reporter construct expression in wild type (wt) and frm larvae. **b**. Frequency of spontaneous swims of wt and frm larvae in dark conditions (i.e., 700nm illumination). Each open circle represents one tracked larva with the number of swim bouts in one minute presented. Also shown are the averages (red circles) ± standard deviation. **c**. Duration of all spontaneous swims in one-minute recording presented in seconds (s) for wt and frm larvae. **d**. Standard deviation (S.D.) of the duration of swim bouts over one minute for each individual larva recorded. **e**. S.D. of the interval between swim bouts over one minute for each individual larva recorded. For **d**. and e., larvae with <4 swim bouts were not included in the analysis. ***, p<0.001; n.s., not significant.

**Figure 8.**
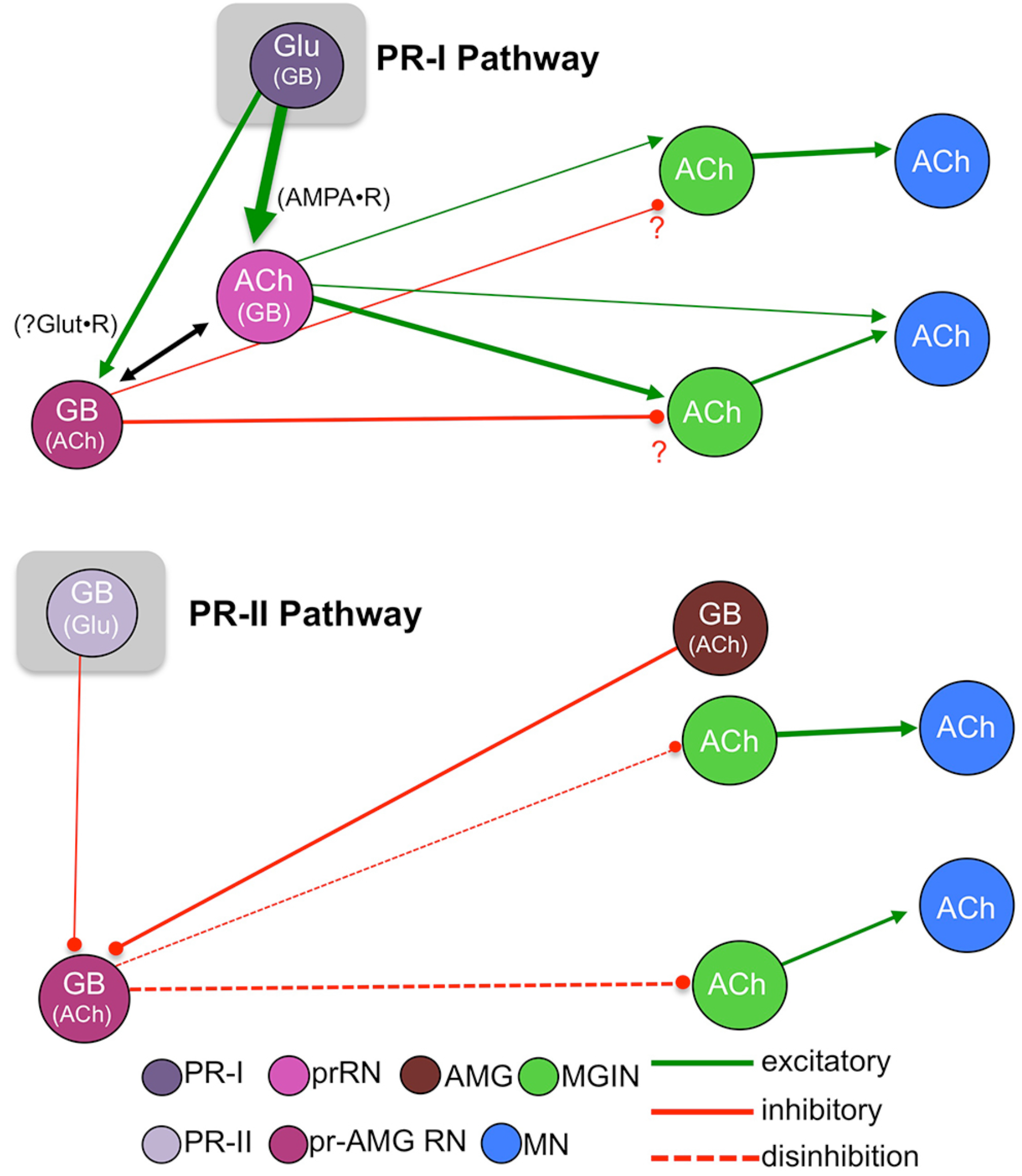
Models showing parallel visuomotor pathways for negative phototaxis (top) and light dimming response (bottom). Neurotransmitters in parentheses are thought to play lesser role in proposed pathway. Abbreviations: PR-II, photoreceptor group II; PR-I photoreceptor group I; pr-AMG RN, photoreceptor ascending motor ganglion relay neuron; prRN, photoreceptor relay neuron; MGIN, motor ganglion interneuron; MN, motor neuron; Glu, glutamate; GB, GABA; ACh, acetylcholine. Cell types are color coded according to [1].

## Discussion

Figure 8 presents a model of the *Ciona* visuomotor circuitry that takes into account the connectome, neurotransmitter use, and behavioral observations. Absent from this model is the detailed and unique connectivity of each neuron in these pathways (sFigure 1a), as well as the inputs from other neurons which are not part of the minimal circuit. Nevertheless, we feel that this model will serve as a useful starting point for more detailed analyses of these components. Our findings support a model for two parallel visuomotor pathways, one mediated by the PR-Is and sensitive to the direction of light, and the other mediated by the PR-IIs and sensitive to changes in ambient light. A number of other sensory systems, including mammalian vision and olfaction and *Drosophila* CO2 detection [40–42] similarly split components of sensory information into parallel circuits. The PR-I circuit is a simple excitatory pathway with glutamatergic photoreceptors projecting to cholinergic prRNs, exciting them via cation-specific ionotropic AMPARs. The prRNs in turn synapse to the cholinergic MGINs, and then these onto the MNs. The fact that glutamate is used by the *Ciona* larvae exclusively in sensory neurons (photoreceptors, antenna cells, and epidermal sensory neurons), coupled with the very limited distribution of AMPARs, allowed us to validate essential components of this circuitry with perampanel. The PR-Is also synapse onto the pr-AMG RNs, which are predicted to be primarily GABAergic. Our observation that AMPAR expression is exclusive to the cholinergic RNs suggests that the response of GABAergic cells to the PR-Is may differ from cholinergic cells, and perhaps plays a role in visual information processing. In fact, the interconnections between the pr-AMG RNs and the AMPAR-expressing prRNs (black arrow Figure 8; see also sFig.1a), are suggestive of an incoherent feedforward loop [43]. We have already documented that *Ciona* larvae are able to phototax in a wide range of illumination conditions [17], and moreover, we have found that *Ciona* larvae show robust fold-change detection [44] behavior (manuscript in preparation). Together these observations suggest that the RN cluster plays a role in visual processing, rather than simply passing information to the MG.

Our model for the PR-II mediated dimming/escape behavior is more surprising, and includes a novelty - inhibitory photoreceptors. From *in situ* hybridization we observed some PR-IIs exclusively express VGAT, while other coexpress VGAT and VGLUT. The significance of VGAT/VGLUT coexpression in the *Ciona* visuomotor pathway is not yet clear, although similar coexpression is widely observed in mammalian brains [45, 46], and invertebrates [47]. It is speculated that co-release of GABA and GLUT may serve to tune excitatory/inhibitory balance. While the connectome shows that not all of the PR-IIs project to the RNs, with a subset instead forming extensive connections to other PR-Is and PR-IIs, the connectome indicates that several of the VGAT-exclusive PR-IIs do project to the pr-AMG RNs (Figure 2e and [1]), consistent with our hypothesis that the PR-II output to the pr-AMG RNs is predominantly inhibitory.

The *Ciona* genome encodes seven ionotropic/Cl^-^ GABA receptor subunit genes (GABAa) [34], but does not have an ortholog of the cationic EXP-1 GABA receptor [48], confirmation that the GABAergic synaptic events are most likely inhibitory. In addition, electrophysiological studies done nearly fifty years ago on larvae of the ascidian *Amaroucium constellatum* reported that their photoreceptors, like those of vertebrates, were hyperpolarizing [49]. In other words, dimming is likely to result in a release of GABA from the PR-IIs. Although the heterogeneity of ascidian photoreceptors (*e.g*., PR-I and -II) was not known at the time, both the vertebrate-like ciliary structure shared by all *Ciona* photoreceptors and the structure of *Ciona* opsins appear to rule out the possibility of depolarizing phototransduction [28, 50]. Also in agreement with an inhibitory output from the PR-IIs is our prediction that the majority of the pr-AMG RNs, the exclusive RNs of the PR-IIs, are themselves GABAergic, which would make a disinhibitory circuit most plausible (Figure 8). We also show that removal of BV sensory input with the *frm* mutant, leads to more frequent spontaneous swims, suggesting that a disinhibitory pathway could lead to stimulation of swimming. Finally, we observed that the AMGs, with the exception of one cell, are GABAergic. The AMGs are one of the primary relay centers for the PNS [12] and project to the MGINs and MNs. However, the AMGs also project ascending axons to the pr-AMG RNs. It is thought that the convergence of PR-II and AMG inputs at the pr-AMG RNs serves to initiate an integrated escape response [12]. Our finding that these two classes of neurons (PR-IIs and AMGs) are likely to have the same input (inhibition) on the pr-AMG RNs further bolsters the integrated response model. Finally, the PR-II mediated dimming response was not inhibited by the AMPAR antagonist perampanel, suggesting the PR-II glutamate release at pr-AMG RNs acts through other receptors, such as the NMDA receptor, and may be more involved in modulating or processing the visual response, and that GABA release may be more important.

Validation of this hypothetical disinhibitory circuit will require analysis of individual neurons in behaving larvae. Although we are able to get robust GCaMP imagery from the CNSs of transgenic *Ciona* larva (our unpublished observations), the fact that the excitation and emission spectra of GCaMP (as well as red-shifted calcium indicators) overlap with the behavioral spectrum of *Ciona*, and the inefficacy of GCaMP for visualizing inhibition, led us to abandon this approach. We are currently exploring methods for electrophysiological recording of *Ciona* BV neurons.

### Ascidians and the evolution of vertebrate visual systems

The evolutionary relationship between the ascidian ocellus and the visual organs of cephalochordates (e.g., amphioxus) and vertebrates remains unclear [50–52]. The observations that the *Ciona* PR-I and PR-II complexes are distinct morphologically, mediate different behaviors, project via distinct visuomotor circuits, and express different NTs, raises the possibility that these two complexes may have independent origins, and thus have different evolutionary relationships to the photoreceptor organs of other chordates. Vertebrates are characterized by the presence of both paired lateral (*i.e.*, retinal) eyes, and an unpaired medial/pineal eye [52]. Amphioxus by contrast, has four distinct photoreceptive organs [53]. The amphioxus *frontal eye* has been proposed as homologous to the vertebrate lateral eyes, while the *lamellar body* is thought to be homologous to the vertebrate pineal organ [53, 54]. The other two amphioxus photoreceptor types, the *dorsal ocelli* and the *Joseph cells*, are thought, based on a number of criteria, including their rhabdomeric morphology-which differs from the ciliary morphology of vertebrate and ascidian photoreceptors-to be vestiges of a more primitive photoreceptive system. Given these relationships, the ascidian PR-I complex is likely to be homologous to the vertebrate lateral eyes and the amphioxus frontal eye, which like the PR-I complex is pigmented and appears to play a role in detecting the direction of light, although not necessarily in taxis [55]. On the other hand, the pineal eyes of amphibian tadpoles and fish larvae mediate a shadow/dimming response, suggesting homology with the ascidian PR-II photoreceptor complex [56, 57]. Nevertheless, the inhibitory nature of the *Ciona* PR-IIs makes assigning homologies more difficult. It is possible that use of GABA by these photoreceptors is a derived feature of ascidians, as inhibitory photoreceptors have yet to be described elsewhere. Alternatively, in the vertebrate retina GABAergic/glycinergic horizontal and amacrine cells are prevalent, and, moreover, it has been proposed that these cells, as well as ganglion cells, are derived from an ancient photoreceptor [51, 58]. While this may imply an alternative evolutionary origin for the *Ciona* PR-IIs, these observations may simply support the plasticity of NT use in visual systems.

## Materials and Methods

### Animals

*Ciona robusta* (a.k.a., *Ciona intestinalis* type A) were either collected from the Santa Barbara Yacht harbor or were obtained from M-REP (San Diego). *Ciona intestinalis* (type B) were obtained from Marine Biological Laboratories (Woods Hole). The mutant*frimousse* (*frm*) and the pVGAT>kaede stable transgenic line (National Bioresource Project, Japan) were cultured at the UC Santa Barbara Marine Lab, as described previously [59]. Larvae were obtained by mixing dissected gametes of 3 adults and cultured in natural seawater at 18°C. Homozygous *frm* larvae were produced by natural spawning of heterozygote *frm* adults.

### Transgene Constructs

*pOpsin1>RFP*. Starting with the plasmid pSP-Ci-opsin 1 (2Kb)>kaede (Takehiro Kusakabe, unpublished), the kaede reading frame was replaced with a synthesized RFP (GeneBlock; IDT).

*pVGAT>H2B::RFP*. The promoter region of VGAT was amplified from genomic DNA using primers containing adaptors for Gateway cloning attB3 and attB5 sites (ataaagtaggctatttaaacaaccagattgcttctgtct and caaaagttgggt tgaggtcgaacgttccg) [25]. This was cloned into pDONR-221-P3-P5 and recombined with an entry clone containing H2B::RFP [60].

### Transgenesis

#### Microinjection

Fertilized one-cell *Ciona intestinalis* (type B) embryos were microinjected through the chorion, as described previously for *C. savignyi* [61].

#### Electroporation

Unfertilized *Ciona robusta* eggs were dechorionated using 0.1% trypsin in 10mM TAPS pH 8.2 in filtered sea water. Eggs were then fertilized and electroporated [62] with 40 μg each of pVACHT>CFP [27] and pVGAT>H2B::RFP. Embryos were cultured at 18°C in filtered sea water with antibiotics until 18 hpf. Larvae were live-mounted for microscopy.

### Hybridization chain reaction (HCR) *in situ*

*Ciona intestinalis*-type B (Marine Biological Laboratories, Woods Hole) were used for *in situ* studies and staged to match the animals used in the connectome study [1]. Optimized HCR *in situ* probes for each target transcript were obtained from Molecular Technologies. For detection of GABAergic/glycinergic cells, probes were made to the vesicular GABA transporter gene (VGAT; accession NM_001032573.1); for glutamatergic cells, probes were made to the vesicular glutamate transporter (VGLUT; NM_001128885.1); for cholinergic cells, probes were made to the vesicular acetylcholine transporter (VACHT; NM_001032789.1). The *in situ* protocol followed the previously published *Ciona in situ* hybridization protocol [63] until the prehybridization step. At this point, the protocol follows the published HCR protocol [30], with the following exception: during the amplification stage, incubation with hairpins is performed for 3 days instead of 12-16 hours.

HCR *in situ* stained larvae were cleared with Slowfade Gold with DAPI (Invitrogen) and imaged on a Leica SP8 resonant scanning confocal microscope. Imaris v. 9.1 [Bitplane] was used to visualize embryos and assign centroids to nuclei using the “add new spots” function, followed by manual correction when necessary. Nuclei were assigned using the maximum intensity projection, cropped to the area of interest.

### anti-GABA staining

*Ciona robusta* embryos were dechorionated at late-tailbud stage as described [63], and then fixed in 4% paraformaldehyde for 24h at 4°C. Fixed larvae were immunostained with a 1:100 dilution of rabbit anti-GABA antibody (Sigma) as described previously [64], and then glycerol cleared and imaged by confocal microscopy.

### Cell Registration

A rotation matrix was calculated based on the 3-dimensional vectors between the anchor cells (ddN and/or antenna cells) and the center of the target cells (photoreceptors or relay neurons) using the HCR *in situ* (target set) and connectome cell centroids (source set). The source set was then rotated to an approximate orientation to the target set. Next, the Coherent Point Drift Algorithm was used to calculate an affine transformation matrix between the source set and the target set of cells [31]. This algorithm models the source set as a Gaussian Mixture Model (GMM), and the target set is treated as observations from the GMM. The transformation matrix is calculated to maximize the Maximum A Posteriori estimation that the observed point cloud is drawn from the GMM. A nearest neighbor mapping based on Euclidean distance is then used to find the closest corresponding point in the target cell set for each cell in the transformed source cell set. The implementation used was adapted from the pure python implementation https://github.com/siavashk/pycpd. The maximum number of iterations was set to 1000 and the maximum root mean squared error for convergence was set to 0.001. The code for the registration is available from Supplemental Materials (Codes 1-3).

#### Confusion matrix

Each dataset containing NT information was registered to every other dataset of the same type using the algorithm detailed above. The EM-registration based cell assignments of each cell in both sets is then compared to each other to see if they agree [32]. The confusion matrix shows the number of times a cell assignment in one dataset corresponds with each other cell assignment in another dataset.

### Behavioral Assays

For time-lapse movies the inverted lid of a 60mm petri dish was first coated with a thin layer of 1% agarose. Larvae were then added to the inverted lid with filtered sea water containing 0.1% BSA with streptomycin and kanamycin each at 20 μg/mL. Finally the dish was covered with a square of glass leaving no air at the top interface. Stock solutions of perampanel were dissolved in methanol and diluted to final concentrations of either 5μm (Santa Cruz Biotech) or 15 μM (Adooq Bioscience) in filtered sea water/BSA/antibiotics. Control samples received methanol alone.

Time-lapse images were collected using a Hamamatsu Orca-ER camera fitted on a Navitar 7000 macro zoom lens. Programmable 700nm and 505nm LED lamps were used to illuminate the larvae (Mightex). All light intensity readings were taken with an Extech Instruments light meter.

#### Dimming-response

All larvae used were between 25 and 28 hours post-fertilization (hpf) (18°C). For image capture, the larvae were illuminated with the 700nm LED lamp and the camera was fitted with a red filter to block the 505nm light. The videos were recorded at 5 fps. In the assays, larvae were first recorded for 10 seconds with the 505nm LED light mounted above the dish at 600 lux and then dimmed to specific values while image capture continued for another 3 minutes. Larvae were allowed to recover for 5 minutes before being assayed again.

#### Phototaxis

All larvae used were approximately 25 hpf (18°C). The 505nm LED light was mounted to one side to the petri dish at approximately 3000 lux. Images were captured at 1 frame per minute for five hours, with the exception of 30 second capture session at 8.9 fps to assay swimming behavior.

#### Spontaneous Swims (frm larvae)

All larvae used were between 26 and 28 hpf. The plates were illuminated with only a 700 nm LED light in order to record dark conditions. The videos were recorded at about 8.9 fps for one minute.

### Behavioral Data analysis

#### Dim-Response criteria

Responses to light dimming were counted if: 1) the larva was stationary at the time of the light dimming, and 2) it swam for longer than 3 seconds. Three seconds was determined by measuring the duration of tail flicks that were previously described [17]. Larvae that bumped or brushed against other larvae or the dish edges were not counted.

#### Tracking and Quantification

Larval swims were tracked using a custom MATLAB script named Estimators of Locomotion Iterations for Animal Experiments (ELIANE). Before uploading to ELIANE, time-lapse images were first processed with Fiji (ImageJ) by subtracting a minimum Z-projection to all the frames and then inverting black and white. ELIANE takes the processed time-lapse images and first creating a background image by averaging the pixels from all the frames. Next, it goes to the initial frame, subtracts the background image, and stores all remaining objects found in the specified region of interest (ROI) as initial objects. Then, analyzing one-by-one the initial objects, it goes frame-by-frame subtracting the background image and analyzing all objects to determine the new position of the object by comparing the Euclidean distances of it to all other objects in that frame. If the object had moved unrealistically fast (>6.5 mm/s), moved outside the ROI, or did not move after a set time (1 minute), the object was not analyzed. This MATLAB script can be found in the Supplemental Materials (Code 4).

The spontaneous swims in the *frimousse* experiment were quantified manually.

#### Sampling

Assessment of larval swim parameters were performed using three independent assays. For the spontaneous swims, which were quantified manually, 25 larvae were selected randomly, starting from the center of the plate going outward, only using the ones that could be tracked for the entire minute recording session.

#### Tests of significance

Dimming response significance and swim frequency were calculated using the Wilcoxon rank-sum test; spontaneous swim time significance was calculated using the Student’ s t-test; and the variance of spontaneous swimming significance was calculated using the F-test.

## Acknowledgments

We thank Takeo Horie and Takahiro Kusakabe for the opsin1 promoter construct; Yasunori Sasakura for the stable pVGAT>kaede line and pVACHT>CFP plasmid; Kerrianne Ryan for her helpful discussion and sharing unpublished data. Chelsea Parlett-Pelleriti for her advice on statistical analysis. We acknowledge the use of the NRI-MCDB Microscopy Facility and the Resonant Scanning Confocal supported by NSF MRI grant 1625770. This work supported by an award from NIH (NS103774) to WCS and BM.

---

**Figure S1.**
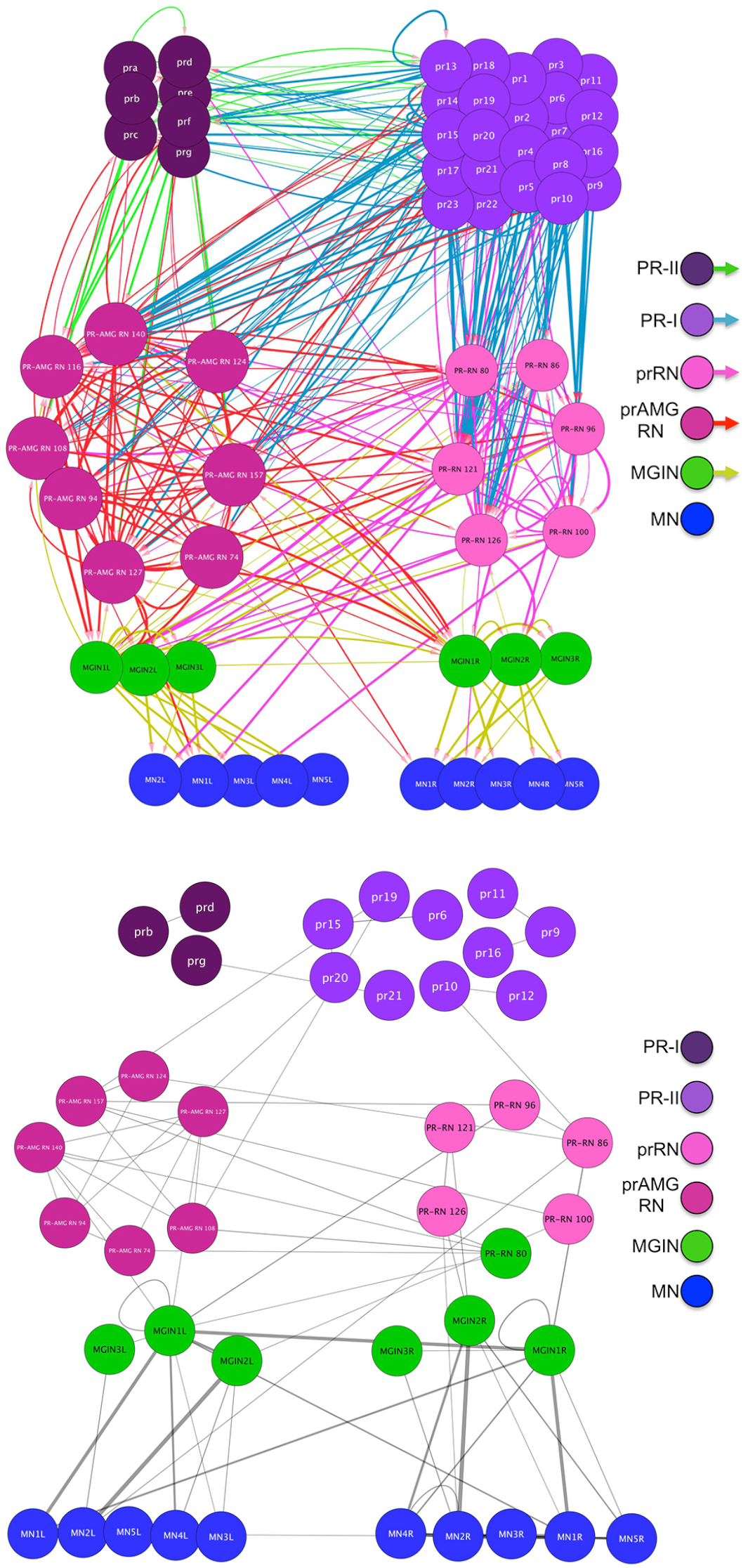
**a. (top)**Chemical synapse connectivity of minimal visuomotor system of *Ciona*. **b. (bottom)**Electrical synapse connectivity of minimal visuomotor system of *Ciona*. Both panels derived from data in [1]. Thickness of lines is proportional to synapse strength. Abbreviations: PR-II, photoreceptor group II; PR-I photoreceptor group I; pr-AMG RN, photoreceptor ascending motor ganglion relay neuron; prRN, photoreceptor relay neuron; MGIN, motor ganglion interneuron. L, left; R, right.

**Figure S2.**
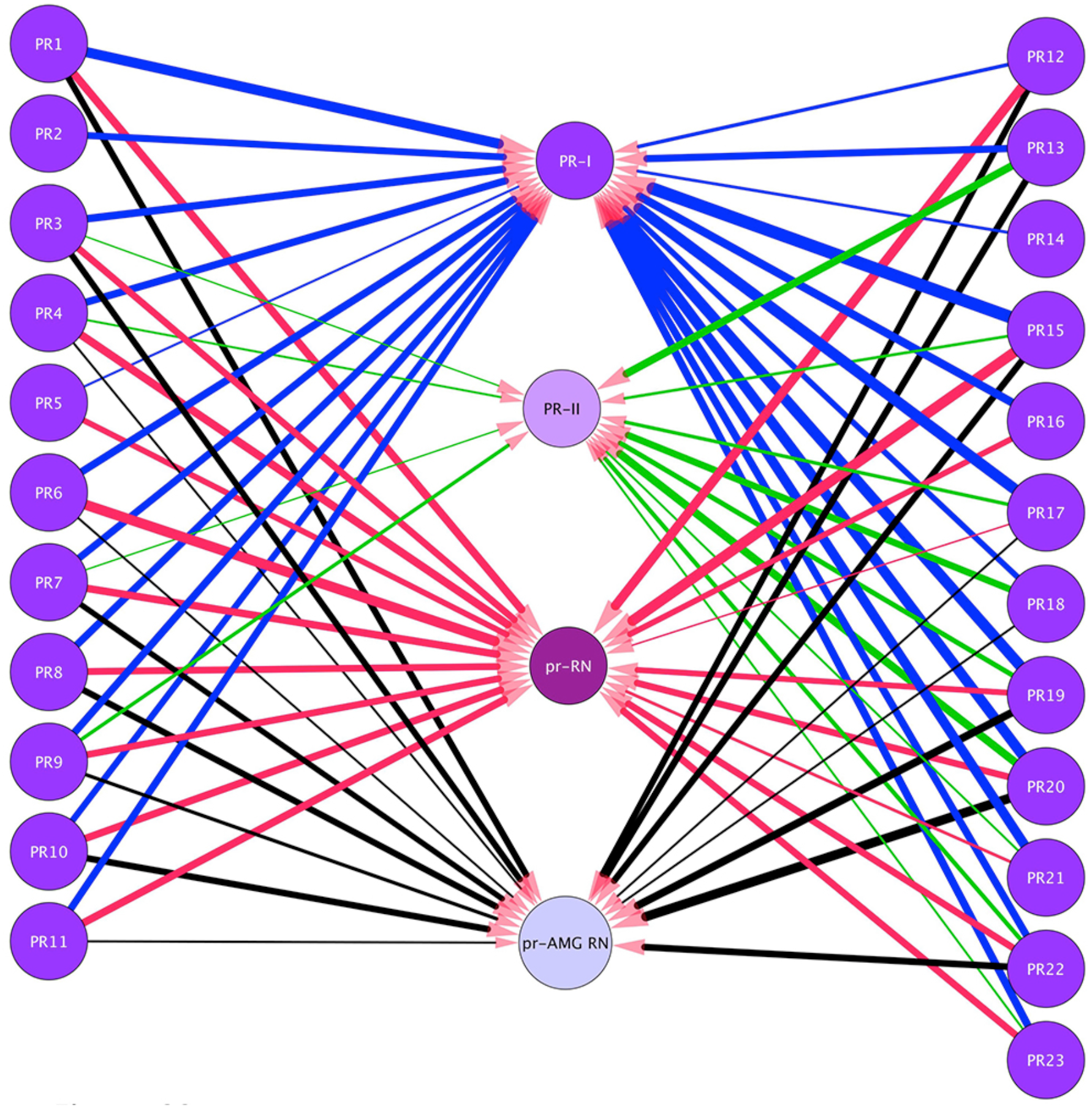
Neurons in the visuomotor circuit postsynaptic to the Group-I Photoreceptors (PR1-PR23). Lines indicate chemical synapses and their thickness indicates synaptic strength. Data from [1]. Abbreviations: PR-I photoreceptor group I; PR-II, photoreceptor group II; pr-AMG RN, photoreceptor ascending motor ganglion relay neuron; prRN, photoreceptor relay neuron.

**Figure S3.**
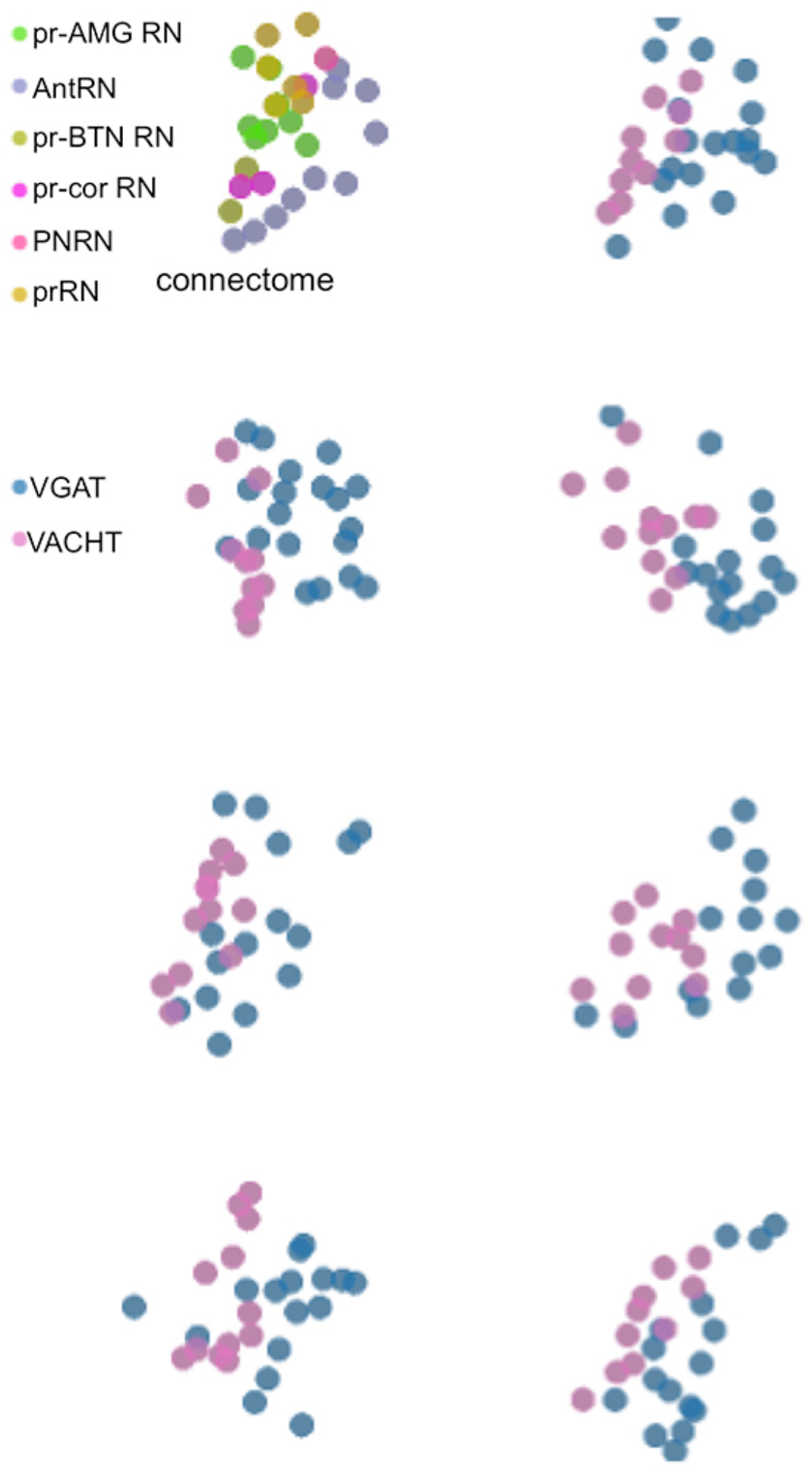
Relay neuron centroids projected in two dimensions. The top left panel is from the connectome, and the remaining panels show centroids from seven larvae. Anterior is to the left and dorsal is at the top.

**Figure S4.**
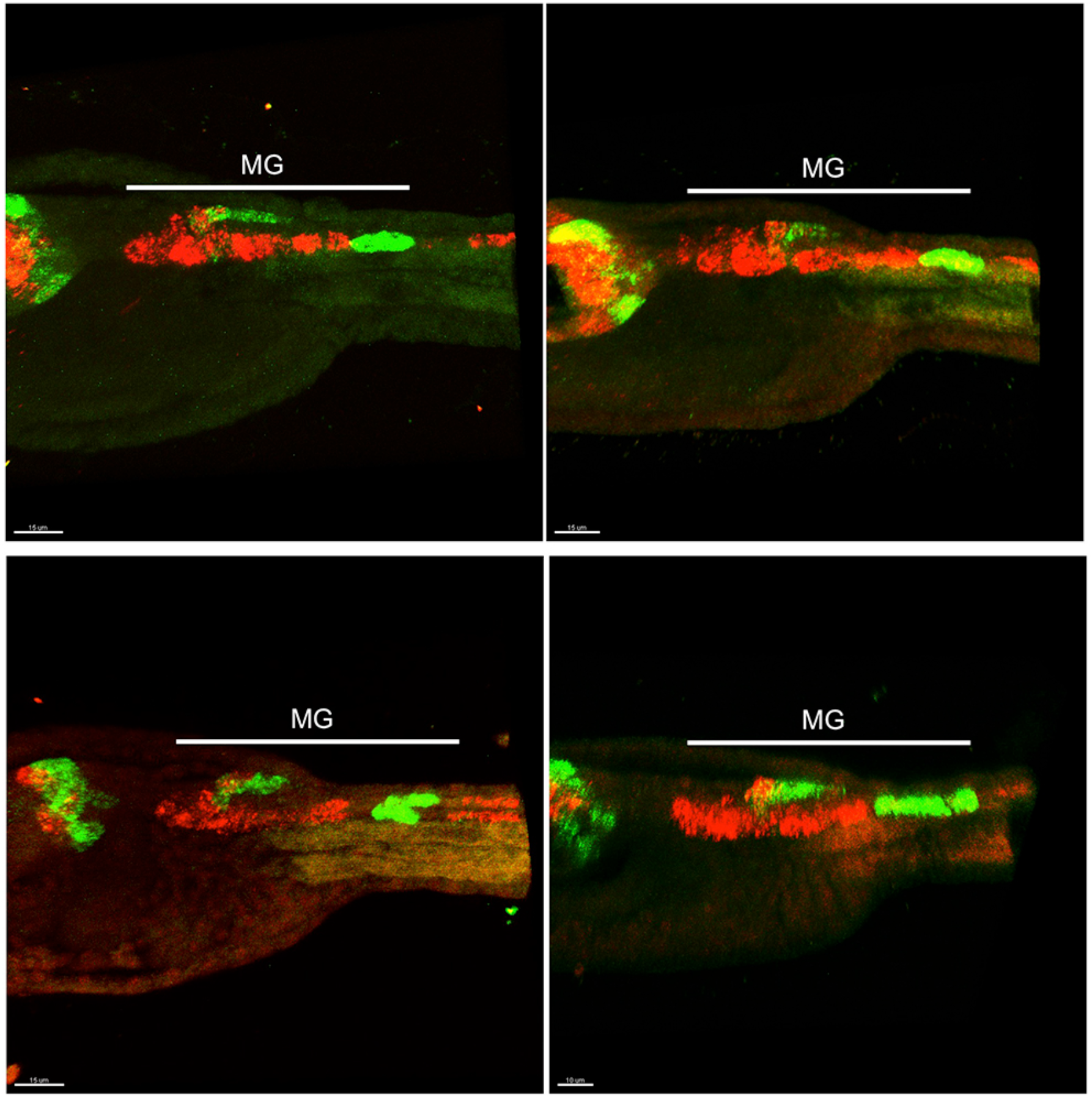
Four representative larvae showing expression pattern for VGAT (green) and VACHT (red) by HCR *in situ*. Anterior is to the left in all sample. MG, motor ganglion.

**Figure S5.**
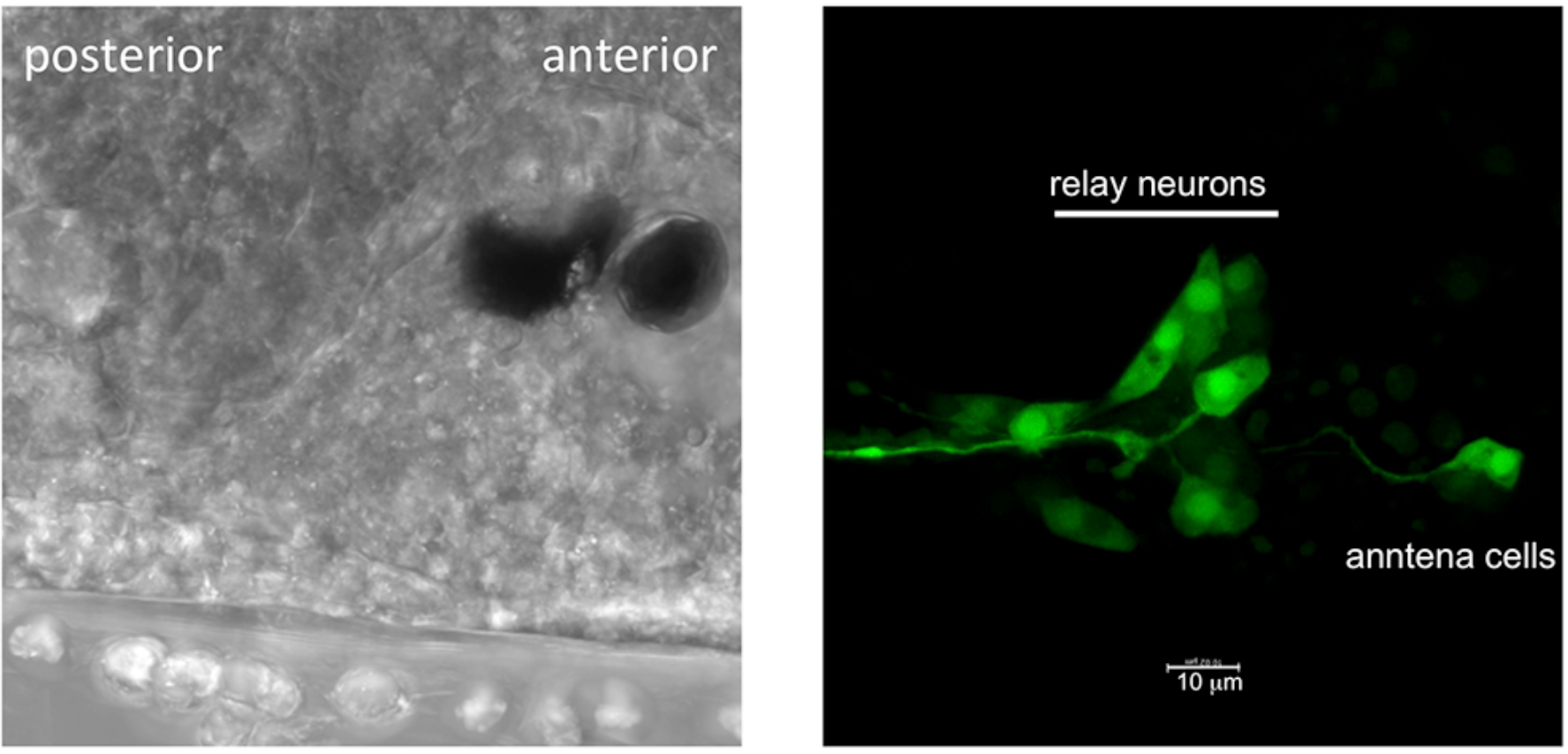
AMPA-receptor neurons in the *Ciona* brain vesicle identified with a AMPA-receptor promoter construct driving GFP.

**Movie1**. Negative phototaxis of control and perampanel-treated *Ciona* larvae in 10cm petri dishes. Directional 505nm illumination is from the left. Frames were taken at 1 per minute over five hours. In the movie the 5 hours is compressed to 15 seconds (*i.e.*, 1200X normal speed). Black and white tones were inverted for the movie to make the larvae more visible.

**Movie 2**. Swimming of control and perampanel-treated *Ciona* larvae in a directional light field. Larvae in 10cm petri dishes were recorded at 9 frames/second. Black and white tones were inverted for the movie to make the larvae more visible. The movie plays at 5X normal speed.

**Movie3**. Dimming response of control and perampanel-treated *Ciona* larvae in 10cm petri dishes. Larvae were imaged for 70 seconds at 9 frames/second, with dimming of 505nm ambient light at 10 seconds. Black and white tones were inverted for the movie, and thus the dimming appears as a brightening. The movie plays at 5X normal speed.

**Movie4**. Spontaneous swimming behavior in wild type and homozygous *frimousse* larvae. Larvae in 10cm petri dishes were recorded for 1 minute at 9 frames/second with 700nm illumination. Black and white tones were inverted for the movie to make the larvae more visible. The movie plays at 5X normal speed.

## References

1. Ryan, K., Z. Lu, and I.A. Meinertzhagen, The CNS connectome of a tadpole larva of Ciona intestinalis (L.) highlights sidedness in the brain of a chordate sibling. Elife, 2016. 5.

2. Randel N., et al., Neuronal connectome of a sensory-motor circuit for visual navigation. Elife, 2014. 3.

3. Eichler K., et al., The complete connectome of a learning and memory centre in an insect brain. Nature, 2017. 548(7666): p. 175–182.

4. Larderet I., et al., Organization of the Drosophila larval visual circuit. Elife, 2017. 6.

5. Helmstaedter M., et al., Connectomic reconstruction of the inner plexiform layer in the mouse retina. Nature, 2013. 500(7461): p. 168–74.

6. Takemura S.Y., et al., A visual motion detection circuit suggested by Drosophila connectomics. Nature, 2013. 500(7461): p. 175–81.

7. Ahrens M.B., et al., Whole-brain functional imaging at cellular resolution using light-sheet microscopy. Nat Methods, 2013. 10(5): p. 413–20.

8. Delsuc F., et al., Tunicates and not cephalochordates are the closest living relatives of vertebrates. Nature, 2006. 439(7079): p. 965–8.

9. Hudson C., The central nervous system of ascidian larvae. Wiley Interdiscip Rev Dev Biol, 2016.

10. Ryan, K., Z. Lu, and I.A. Meinertzhagen, Circuit Homology between Decussating Pathways in the Ciona Larval CNS and the Vertebrate Startle-Response Pathway. Curr Biol, 2017. 27(5): p. 721–728.

11. Imai J.H. and I.A. Meinertzhagen, Neurons of the ascidian larval nervous system in Ciona intestinalis: II. Peripheral nervous system. J Comp Neurol, 2007. 501(3): p. 335–52.

12. Ryan, K., Z. Lu, and I.A. Meinertzhagen, The peripheral nervous system of the ascidian tadpole larva: Types of neurons and their synaptic networks. J Comp Neurol, 2018. 526(4): p. 583–608.

13. Stolfi A., et al., Migratory neuronal progenitors arise from the neural plate borders in tunicates. Nature, 2015. 527(7578): p. 371–4.

14. Lemaire P., Evolutionary crossroads in developmental biology: the tunicates. Development, 2011. 138(11): p. 2143–52.

15. Kajiwara S. and M. Yoshida, Changes in Behavior and Ocellar Structure during the Larval Life of Solitary Ascidians. Biological Bulletin, 1985. 169(3): p. 565–577.

16. Sakurai D., et al., The role of pigment cells in the brain of ascidian larva. J Comp Neurol, 2004. 475(1): p. 70–82.

17. Salas P., et al., Photoreceptor specialization and the visuomotor repertoire of the primitive chordate Ciona. J Exp Biol, 2018. 221 (Pt 7).

18. Svane I.B. and C.M. Young, The ecology and behavior of ascidian larvae. Mar Biol Rev, 1989. 27(45-90).

19. Moret F., et al., The dopamine-synthesizing cells in the swimming larva of the tunicate Ciona intestinalis are located only in the hypothalamus-related domain of the sensory vesicle. Eur J Neurosci, 2005. 21(11): p. 3043–55.

20. Horie T., et al., Pigmented and nonpigmented ocelli in the brain vesicle of the ascidian larva. J Comp Neurol, 2008. 509(1): p. 88–102.

21. Brown E.R., et al., GABAergic synaptic transmission modulates swimming in the ascidian larva. Eur J Neurosci, 2005. 22(10): p. 2541–8.

22. Horie, T., T. Kusakabe, and M. Tsuda, Glutamatergic networks in the Ciona intestinalis larva. J Comp Neurol, 2008. 508(2): p. 249–63.

23. Pennati R., et al., Developmental expression of tryptophan hydroxylase gene in Ciona intestinalis. Dev Genes Evol, 2007. 217(4): p. 307–13.

24. Takamura, K., N. Minamida, and S. Okabe, Neural map of the larval central nervous system in the ascidian Ciona intestinalis. Zoolog Sci, 2010. 27(2): p. 191–203.

25. Yoshida R., et al., Identification of neuron-specific promoters in Ciona intestinalis. Genesis, 2004. 39(2): p. 130–40.

26. Zega G., et al., Developmental expression of glutamic acid decarboxylase and of gamma-aminobutyric acid type B receptors in the ascidian Ciona intestinalis. J Comp Neurol, 2008. 506(3): p. 489–505.

27. Horie T., et al., Ependymal cells of chordate larvae are stem-like cells that form the adult nervous system. Nature, 2011. 469(7331): p. 525–8.

28. Kusakabe T., et al., Ci-opsin1, a vertebrate-type opsin gene, expressed in the larval ocellus of the ascidian Ciona intestinalis. FEbS Lett, 2001. 506(1): p. 69–72.

29. Kusakabe T., et al., Computational discovery of DNA motifs associated with cell type-specific gene expression in Ciona. Dev Biol, 2004. 276(2): p. 563–80.

30. Choi H.M.T., et al., Third-generation in situ hybridization chain reaction: multiplexed, quantitative, sensitive, versatile, robust. Development, 2018. 145(12).

31. Myronenko A. and X. Song, Point set registration: coherent point drift. IEEE Trans Pattern Anal Mach Intell, 2010. 32(12): p. 2262–75.

32. Stehman S.V., Selecting and interpreting measures of thematic classification accuracy. Remote Sensing of Environment, 1997. 62(1): p. 77–89.

33. Takamura K., et al., Developmental expression of ascidian neurotransmitter synthesis genes. I. Choline acetyltransferase and acetylcholine transporter genes. Dev Genes Evol, 2002. 212(1): p. 50–3.

34. Okamura Y., et al., Comprehensive analysis of the ascidian genome reveals novel insights into the molecular evolution of ion channel genes. Physiol Genomics, 2005. 22(3): p. 269–82.

35. Hirai S., et al., AMPA glutamate receptors are required for sensory-organ formation and morphogenesis in the basal chordate. Proc Natl Acad Sci U S A, 2017. 114(15): p. 3939–3944.

36. Hanada T., et al., Perampanel: a novel, orally active, noncompetitive AMPA-receptor antagonist that reduces seizure activity in rodent models of epilepsy. Epilepsia, 2011. 52(7): p. 1331–40.

37. Deschet K. and W.C. Smith Frimousse — a spontaneous ascidian mutant with anterior ectodermal fate transformation. Current Biology, 2004. 14: p. R408-R410.

38. Hackley C., et al., A transiently expressed connexin is essential for anterior neural plate development in Ciona intestinalis. Development, 2013. 140(1): p. 147–55.

39. Nakagawa M., et al., Action spectrum for the photophobic response of Ciona intestinalis (Ascidieacea, Urochordata) larvae implicates retinal protein. Photochem Photobiol, 1999. 70(3): p. 359–62.

40. Callaway E.M., Structure and function of parallel pathways in the primate early visual system. J Physiol, 2005. 566(Pt 1): p. 13–9.

41. Geramita, M.A., S.D. Burton, and N.N. Urban, Distinct lateral inhibitory circuits drive parallel processing of sensory information in the mammalian olfactory bulb. Elife, 2016. 5.

42. Lin H.H., et al., Parallel neural pathways mediate CO2 avoidance responses in Drosophila. Science, 2013. 340(6138): p. 1338–41.

43. Alon U., Network motifs: theory and experimental approaches. Nat Rev Genet, 2007. 8(6): p. 450–61.

44. Adler M. and U. Alon, Fold-change detection in biological systems. Current Opinion in Systems Biology, 2018. 8: p. 81-89.

45. Fattorini G., et al., Co-expression of VGLUT1 and VGAT sustains glutamate and GABA co-release and is regulated by activity in cortical neurons. J Cell Sci, 2015. 128(9): p. 1669–73.

46. Zander J.F., et al., Synaptic and vesicular coexistence of VGLUT and VGAT in selected excitatory and inhibitory synapses. J Neurosci, 2010. 30(22): p. 7634–45.

47. Fabian-Fine R., et al., Co-localization of Gamma-Aminobutyric Acid and Glutamate in Neurons of the Spider Central Nervous System. Cell Tissue Res, 2015. 362(3): p. 461-79.

48. Beg A.A. and E.M. Jorgensen, EXP-1 is an excitatory GABA-gated cation channel. Nat Neurosci, 2003. 6(11): p. 1145–52.

49. Gorman, A.L., J.S. McReynolds, and S.N. Barnes, Photoreceptors in primitive chordates: fine structure, hyperpolarizing receptor potentials, and evolution. Science, 1971. 172(3987): p. 1052–4.

50. Kusakabe T. and M. Tsuda, Photoreceptive systems in ascidians. Photochem Photobiol, 2007. 83(2): p. 248–52.

51. Lamb T.D., Evolution of phototransduction, vertebrate photoreceptors and retina. Prog Retin Eye Res, 2013. 36: p. 52-119.

52. Lamb, T.D., S.P. Collin, and E.N. Pugh, Jr., Evolution of the vertebrate eye: opsins, photoreceptors, retina and eye cup. Nat Rev Neurosci, 2007. 8(12): p. 960–76.

53. Pergner J. and Z. Kozmik, Amphioxus photoreceptors - insights into the evolution of vertebrate opsins, vision and circadian rhythmicity. Int J Dev Biol, 2017. 61(10-11-12): p. 665–681.

54. Vopalensky P., et al., Molecular analysis of the amphioxus frontal eye unravels the evolutionary origin of the retina and pigment cells of the vertebrate eye. Proc Natl Acad Sci U S A, 2012. 109(38): p. 15383–8.

55. Stokes M.D. and N.D. Holland, Ciliary Hovering in Larval Lancelets (=Amphioxus). Biol Bull, 1995. 188(3): p. 231–233.

56. Jamieson D. and A. Roberts, Responses of young Xenopus laevis tadpoles to light dimming: possible roles for the pineal eye. J Exp Biol, 2000. 203(Pt 12): p. 1857–67.

57. Yoshizawa M. and W.R. Jeffery, Shadow response in the blind cavefish Astyanax reveals conservation of a functional pineal eye. J Exp Biol, 2008. 211 (Pt 3): p. 292–9.

58. Arendt D., Evolution of eyes and photoreceptor cell types. Int J Dev Biol, 2003. 47(7-8): p. 563–71.

59. Veeman, M.T., S. Chiba, and W.C. Smith, Ciona genetics. Methods Mol Biol, 2011. 770: p. 401-22.

60. Roure A., et al., A multicassette Gateway vector set for high throughput and comparative analyses in ciona and vertebrate embryos. PLoS One, 2007. 2(9): p. e916.

61. Deschet, K., Y. Nakatani, and W.C. Smith, Generation of Ci-Brachyury-GFP stable transgenic lines in the ascidian Ciona savignyi. Genesis, 2003. 35(4): p. 248–59.

62. Zeller R.W., Electroporation in Ascidians: History, Theory and Protocols. Adv Exp Med Biol, 2018. 1029: p. 37-48.

63. Corbo, J.C., M. Levine, and R.W. Zeller, Characterization of a notochord-specific enhancer from the Brachyury promoter region of the ascidian, Ciona intestinalis. Development, 1997. 124(3): p. 589–602.

64. Veeman M.T., et al., Chongmague reveals an essential role for laminin-mediated boundary formation in chordate convergence and extension movements. Development, 2008. 135(1): p. 33–41.

